# TrkB-expressing paraventricular hypothalamic neurons suppress appetite through multiple neurocircuits

**DOI:** 10.1101/671875

**Authors:** Juan Ji An, Clint E. Kinney, Guey-Ying Liao, Eric J. Kremer, Baoji Xu

## Abstract

The TrkB receptor is critical for the control of energy balance, as mutations in its gene (*NTRK2*) lead to hyperphagia and severe obesity in humans and mice. The main neural substrate mediating the appetite-suppressing activity of TrkB, however, remains unknown. Here, we demonstrate that selective *Ntrk2* deletion within the paraventricular hypothalamus (PVH) leads to severe hyperphagic obesity. Furthermore, chemogenetic activation or inhibition of TrkB-expressing PVH (PVH^TrkB^) neurons suppresses or increases food intake, respectively. PVH^TrkB^ neurons project to multiple brain regions, including the ventromedial hypothalamus (VMH) and the lateral parabrachial nucleus (LPBN). We found that PVH^TrkB^ neurons projecting to LPBN are distinct from those projecting to VMH, yet *Ntrk2* deletion in PVH neurons projecting to either VMH or LPBN results in hyperphagia and obesity. Therefore, TrkB signaling is a key regulator of a previously uncharacterized and heterogenous neuronal population within the PVH that impinges upon multiple circuits to govern appetite.

## INTRODUCTION

The regulation of food intake is essential for survival in all organisms, and its dysregulation not only affects survival but also causes health problems such as obesity. Several ligand-receptor pairs play a crucial role in the central regulation of energy balance. They include leptin and the leptin receptor, alpha-melanocyte-stimulating hormone (αMSH) and the melanocortin-4 receptor (MC4R), and brain-derived neurotrophic factor (BDNF) and the TrkB receptor. Defects in any of these three ligand-receptor pairs lead to severe obesity in humans^1–5^ and mice^6–11^. Furthermore, several common single nucleotide polymorphisms in the *MC4R* and *BDNF* genes have been found to be associated with increased body mass index in large-scale genome-wide association studies^12–15^. Remarkable progress has been made in elucidation of neural circuits that mediate the effects of leptin and αMSH on energy balance^16^. By comparison, much less is known about the neural substrate through which the BDNF-TrkB pathway regulates appetite and energy expenditure.

BDNF is well known for its role in neuronal development and synaptic function^17–19^. It signals through the binding of two distinct classes of receptor proteins: the tropomyosin receptor kinase B (TrkB) and the p75 neurotrophin receptor (p75^NTR^). Mutations in the *NTRK2* gene, which codes for TrkB, lead to severe obesity in mice and humans^4,11^. Conversely, no obesity phenotype has been observed in mice lacking the p75^NTR^ receptor^20^. In fact, mice lacking the p75^NTR^ receptor are protected from obesity induced by high-fat diet and remain lean due to increased energy expenditure^21^. Therefore, BDNF should signal through TrkB to regulate energy balance. The paraventricular hypothalamus (PVH) and the ventromedial hypothalamus (VMH) have been identified as two important brain areas that express BDNF to suppress food intake and promote energy expenditure^22–24^. However, it remains unclear in which population of hypothalamic neurons TrkB is required for the control of energy balance and how these TrkB-expressing neurons regulate food intake and/or energy expenditure. Our recent work found that *Ntrk2* deletion in the dorsomedial hypothalamus (DMH) led to modest hyperphagia and obesity^26^, but to a much lesser extent in comparison with *Ntrk2* hypomorphic mice in which *Ntrk2* expression is reduced to a quarter of the normal amount throughout the body^11^, indicating that the main neural substrate underlying the action of TrkB signaling on energy balance remains to be identified.

Here we report that TrkB-expressing PVH (PVH^TrkB^) neurons are distinct from neurons that express BDNF, MC4R, or prodynorphin (PDYN) in the PVH, the three subtypes of neurons that are known to suppress appetite^22,27,28^. We found that chemogenetic inhibition of PVH^TrkB^ neurons greatly increased food intake, whereas chemogenetic activation of these neurons drastically reduced food intake. Furthermore, we found that deletion of the *Ntrk2* gene in the PVH led to severe hyperphagic obesity. We then designed a projection-specific gene deletion approach and used this approach to reveal that PVH^TrkB^ neurons project to the VMH and the lateral parabrachial nucleus (LPBN) to suppress food intake. These studies not only identify the main neural substrate by which TrkB signaling regulates energy balance, but also implicates the PVH → VMH neurocircuit in the control of appetite.

## RESULTS

### Many TrkB neurons in the PVH are distinct from previously defined ones

The PVH is a heterogeneous brain structure with many different cell types^29,30^. At least 10 types of PVH neurons can be molecularly defined by the expression of BDNF, corticotropin-releasing hormone (CRH), growth hormone-releasing hormone (GHRH), MC4R, oxytocin, prodynorphin (PDYN), somatostatin, tyrosine hydroxylase, thyrotropin-releasing hormone (TRH), and vasopressin^22,27,28,31^. We previously found that approximately 5% PVH^TrkB^ neurons expressed BDNF^22^. We sought to determine whether PVH^TrkB^ neurons belong to other defined neuronal populations.

TrkB antibodies do not mark cell bodies well in brain sections, because TrkB is distributed to the membrane of both cell bodies and processes. We employed the *Ntrk2*^*CreER*/+^ mouse strain^32^, in which the Cre-ERT2 sequence is inserted into the *Ntrk2* locus immediately after the first coding ATG, for colocalization studies. We crossed *Ntrk2*^*CreER*/+^ mice to *Rosa26*^Ai9/+^ mice^33^ to generate *Ntrk2*^*CreER*/+^*;Rosa26*^*Ai9*/+^ mice, in which TrkB-expressing cells express tdTomato after tamoxifen treatment. Since the kinase-lacking truncated TrkB is expressed in astrocytes^34^, tdTomato should also label astrocytes. We detected many tdTomato-labelled TrkB cells in the PVH (Fig. 1a), which are either neurons that are positive for the neuronal marker NeuN (Supplementary Fig. 1a) or astrocytes that are positive for glial fibrillary acidic protein (Supplementary Fig. 1b). We noticed that there was a high colocalization between TrkB and oxytocin in the PVH (18% TrkB neurons express oxytocin and 51% oxytocin neurons express TrkB; Fig. 1b). In addition, a small number of PVH^TrkB^ neurons express TRH (2% TrkB neurons express TRH and 12% TRH neurons express TrkB; Fig. 1c), PDYN (6% TrkB neurons express PDYN and 17% PDYN neurons express TrkB; Fig. 1d), somatostatin (2% TrkB neurons express somatostatin and 16% somatostatin neurons express TrkB; Supplementary Fig. 1c) and GHRH (5% TrkB neurons express GHRH and 30% GHRH neurons express TrkB; Supplementary Fig. 1d). Note that higher percentages of PVH^TrkB^ neurons are positive for these markers in the shown images (Fig. 1 and Supplementary Fig. 1), because the images were taken from the area where these markers are expressed. Very few PVH^TrkB^ neurons express CRH (Fig. 1e), tyrosine hydroxylase (Supplementary Fig. 1e), or vasopressin (Supplementary Fig. 1f). We also examined colocalization of TrkB with MC4R in the PVH using the BAC *Mc4r-EGFP* mouse strain^35^. We only detected low colocalization of TrkB with MC4R in the PVH of tamoxifen-treated *Ntrk2^CreER/+^;Rosa26^Ai9/+^;Mc4r-EGFP* mice (3% TrkB neurons express MC4R and 10% MC4R neurons express TrkB; Fig. 1f). These results indicate that the majority of PVH^TrkB^ neurons are distinct from previously defined neuronal populations.

**Figure 1.**
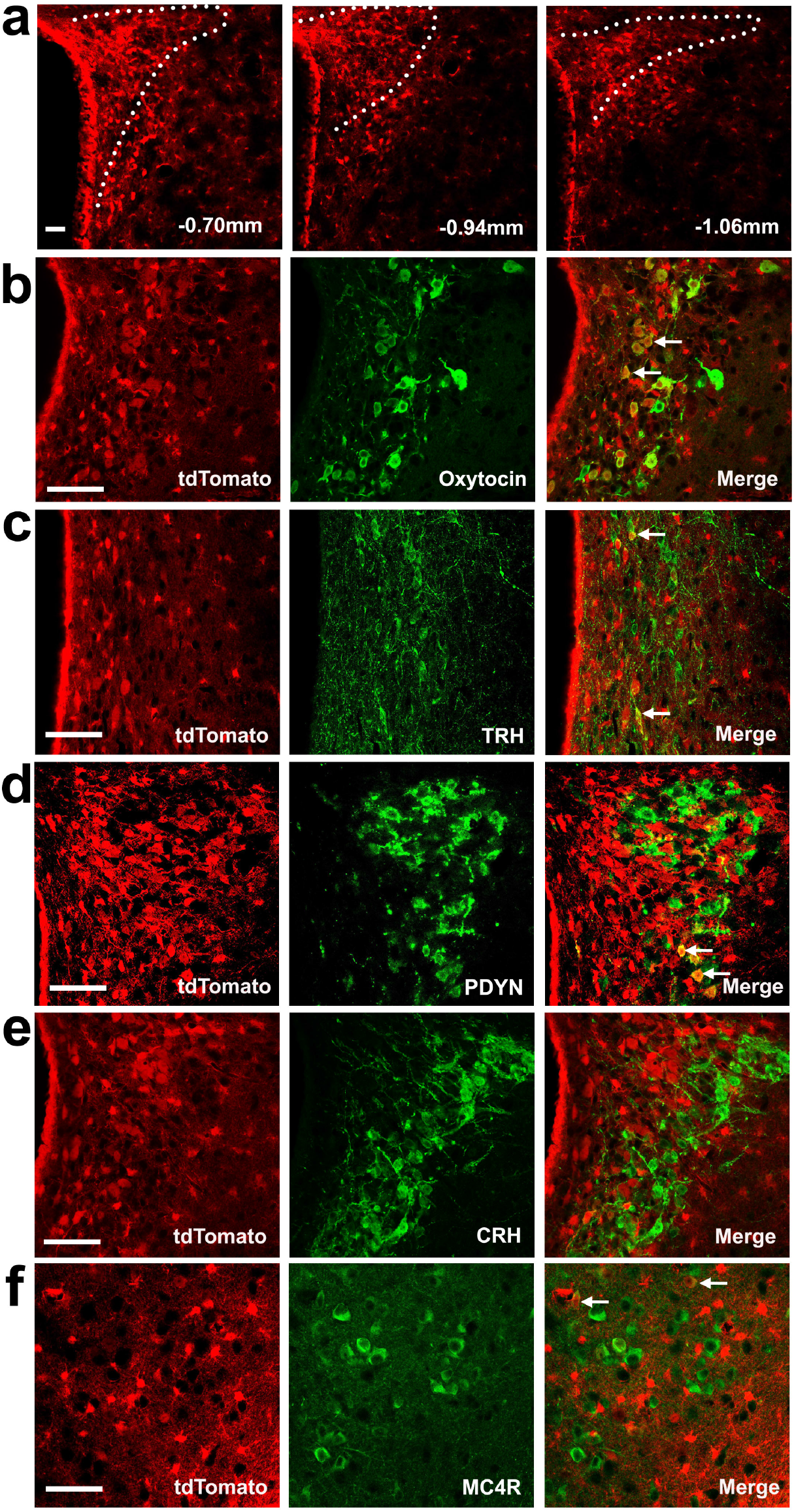
Co-expression of TrkB with other neuronal markers in the PVH. TrkB-expressing cells are marked by tdTomato in tamoxifen-treated *Ntrk2*^*CreER*/+^*;Rosa26*^*Ai9*/+^ mice. (**a**) Distribution of TrkB-expressing cells in the PVH at 3 rostral-caudal positions. (**b-e**) Co-expression of TrkB with oxytocin, TRH, PDYN and CRH. (**f**) Co-expression of tdTomato and EGFP in *Ntrk2*^*CreER*/+^*;Rosa26*^*Ai9*/+^*;Mc4r-EGFP* mice. Arrows denote representative neurons expressing both TrkB and a PVH marker. Scale bars are 50 μm long.

### Deletion of the *Ntrk2* gene with Sim1-Cre leads to obesity

We previously reported that *Ntrk2* deletion in several hypothalamic areas using the Rgs9-Cre led to obesity^25^; however, the obesity phenotype was not as severe as what we observed in *Ntrk2* hypomorphic mice that express TrkB at ~25% of the normal amount throughout the body^11,25^. Given that Rgs9-Cre only deletes *Ntrk2* in a small number of PVH neurons^25^ and that stereotaxic injection of recombinant BDNF into the PVH reduces food intake and increases energy expenditure^36,37^, this observation raises the possibility that TrkB expressed in the PVH (PVH TrkB hereafter) plays an important role in the control of energy balance. We tested this possibility by deleting *Ntrk2* in the PVH.

We tested whether Sim1-Cre^38^ is suitable for abolishment of *Ntrk2* gene expression in the PVH by crossing Sim1-Cre mice to *Ntrk2*^*fBZ*/+^ mice to generate Sim1-Cre*;Ntrk2*^*fBZ*/+^ mice, in which β-galactosidase is expressed in otherwise TrkB-expressing neurons once the floxed *Ntrk2*^*fBZ*^ allele is deleted by Cre recombinase as shown in our previous studies^39,40^. We detected many cells expressing β-galactosidase in the PVH (Fig. 2a), suggesting that Sim1-Cre is effective in abolishing TrkB expression in the PVH. In addition to the PVH, Sim1-Cre can also abolish TrkB expression in some neurons of other brain regions including the cerebral cortex, hippocampus, amygdala, thalamus and DMH (Supplementary Fig. 2a-f).

**Figure 2.**
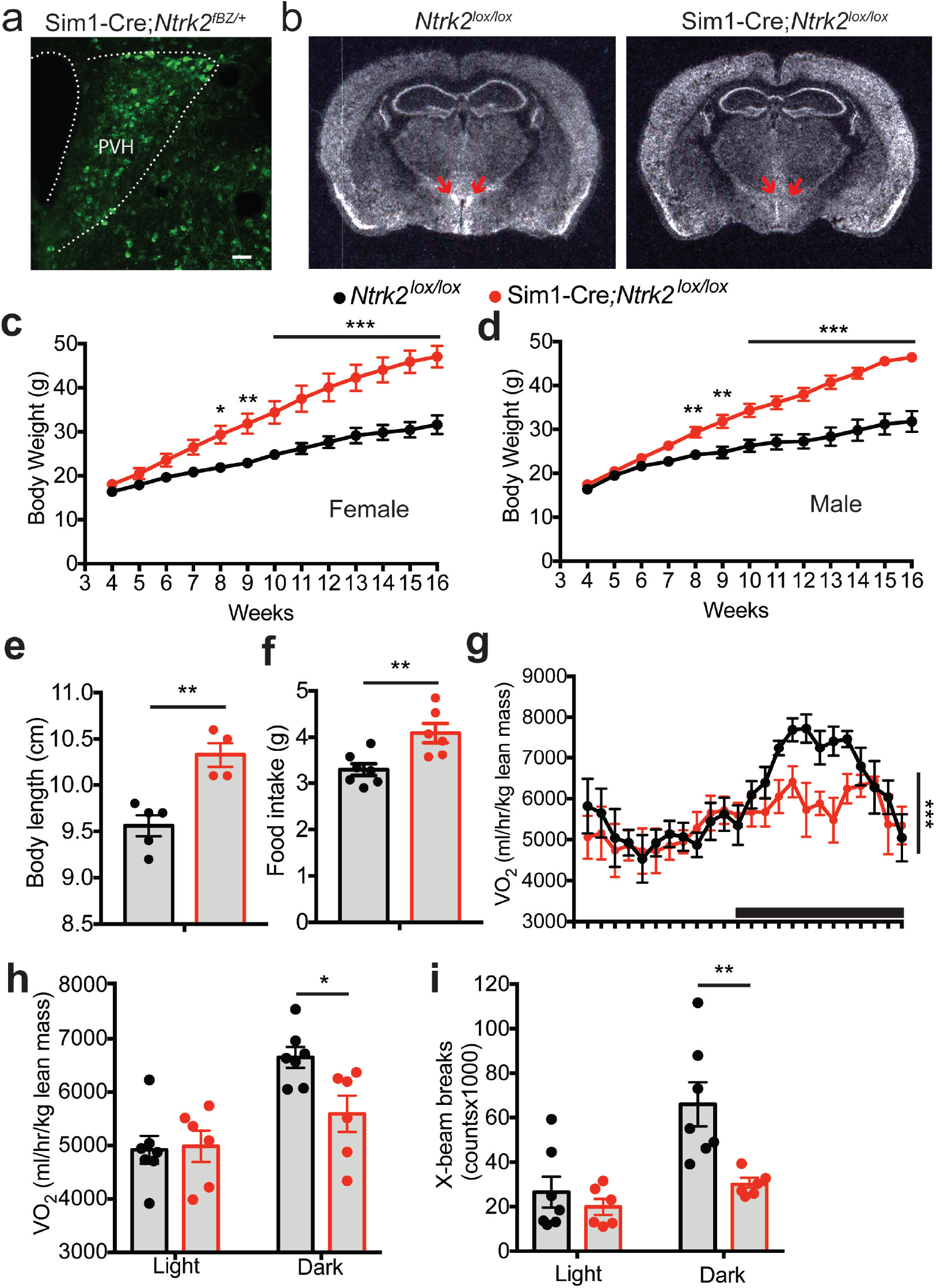
Deletion of the *Ntrk2* gene in Sim1-Cre cells led to obesity. (**a**) β-galactosidase immunohistochemistry of Sim1-Cre*;Ntrk2*^*fBZ*/+^ mice, showing *Ntrk2* deletion in the PVH. Scale bar, 50 μm. (**b**) *In situ* hybridization of brain sections from 4-month-old *Ntrk2*^*lox/lox*^ mice (control) and Sim1-Cre*;Ntrk2*^*lox/lox*^ mice (mutant) reveals that *Ntrk2* gene expression was abolished in the PVH of mutant mice. Arrows denote the PVH. (**c**) Body weight of female control and mutant mice. n = 5 mice per genotype. Two-way ANOVA with post hoc Bonferroni multiple comparisons; *F*(1, 104) = 163.7, *P* < 0.0001 for genotype; \**P* < 0.05, \*\**P* < 0.01, and \*\*\**P* < 0.001. (**d**) Body weight of male control and mutant mice. n = 4 controls and 6 mutants. Two-way ANOVA with post hoc Bonferroni multiple comparisons; *F*(1, 104) = 193.5, *P* < 0.0001 for genotype; \*\**P* < 0.01 and \*\*\**P* < 0.001. (**e**) Body length of 8-wk-old female control and mutant mice. n = 5 controls and 4 mutants. Student’s *t* test; \*\**P* < 0.01. (**f**) Daily food intake of 8-wk-old female control and mutant mice. n = 7 controls and 6 mutants. Student’s *t* test; \*\**P* < 0.01. (**g**) Distribution of oxygen consumption (VO_2_) over a 24-h period in 6-wk-old female mice. The black bar above *x* axis indicates the dark cycle. n = 7 controls and 6 mutants. Two-way ANOVA for genotype; *F*(1, 264) = 14.54, \*\*\**P* = 0.0002. (**h**) VO_2_ of 6-wk-old female mice during the light and dark cycles. n = 7 controls and 6 mutants. Student’s *t* test; \*\**P* < 0.01. (**D**) Locomotor activity of 6-wk-old female mice during the light and dark cycles. n = 7 controls and 6 mutants. Student’s *t* test; \*\**P* < 0.01. Error bars indicate SEM.

We crossed Sim1-Cre mice to *Ntrk2*^*lox/lox*^ mice, which express a normal amount of TrkB in the absence of Cre^41^, to produce *Ntrk2*^*lox/lox*^ (control) and Sim1-Cre*;Ntrk2*^*lox/lox*^ (mutant) mice. *In situ* hybridization confirmed that *Ntrk2* expression was abolished in the PVH of Sim1-Cre*;Ntrk2*^*lox/lox*^ mice (Fig. 2b). Male and female Sim1-Cre*;Ntrk2*^*lox/lox*^ mice at 16 weeks of age were 45% and 49% heavier than sex-matched control mice, respectively (Fig. 2c, d). The Sim1-Cre*;Ntrk2*^*lox/lox*^ mice also had a longer body (Fig. 2e) and larger adipose tissues relative to control mice (Supplementary Fig 2g, h). These results indicate that Sim1-Cre*;Ntrk2*^*lox/lox*^ mice develop obesity.

We then determined if increased energy intake and/or reduced energy expenditure contribute to the obesity phenotype displayed by Sim1-Cre*;Ntrk2*^*lox/lox*^ mice. At 8 weeks of age, female Sim1-Cre*;Ntrk2*^*lox/lox*^ mice daily consumed 24% more food than control mice (Fig. 2f). We employed indirect calorimetry to estimate energy expenditure in mice at 6 weeks of age when control and Sim1-Cre*;Ntrk2*^*lox/lox*^ mice had comparable body weights (Fig 2c). Control and Sim1-Cre*;Ntrk2*^*lox/lox*^ mice had comparable oxygen consumption (VO_2_) during the light period when mice are mostly at sleep (Fig. 2g, h), suggesting Sim1-Cre*;Ntrk2*^*lox/lox*^ mice have a relatively normal basal metabolic rate. During the dark period, VO_2_ and locomotor activity in Sim1-Cre*;Ntrk2*^*lox/lox*^ mice were reduced by 15% and 51%, respectively, compared with control mice (Fig. 2g-i). These results indicate that TrkB signaling in Sim1-Cre cells promote physical activity and energy expenditure. Collectively, our results indicate that Sim1-Cre*;Ntrk2*^*lox/lox*^ mice develop obesity due to increased food intake and reduced energy expenditure.

### Selective *Ntrk2* deletion in the adult PVH led to hyperphagic obesity

Sim1-Cre is expressed in other brain regions in addition to the PVH^38^ (Supplementary Fig. 2a-f), so the obesity phenotype in Sim1-Cre*;Ntrk2*^*lox/lox*^ mice might result from *Ntrk2* deletion in non-PVH neurons or cells. To validate a role for PVH TrkB in the regulation of energy balance, we stereotaxically injected neuron-tropic AAV2, either AAV2-CMV-Cre-GFP or AAV2-CMV-GFP, into the PVH of 8-wk-old female *Ntrk2*^*lox/lox*^ mice (Fig. 3a). Injection of AAV2-CMV-Cre-GFP selectively abolished *Ntrk2* expression in the PVH (Fig. 3a2). Mice injected with AAV2-CMV-Cre-GFP are divided into two groups based on the extent of AAV infection in the PVH: AAV2-CMV-Cre-GFP (missed) and AAV2-CMV-Cre-GFP (hit) (Supplementary Table 1). Mice in the AAV2-CMV-Cre-GFP (hit) group developed severe obesity quickly, compared to mice in the AAV2-CMV-GFP group and the AAV2-CMV-Cre-GFP (Missed) group (Fig. 3b, c). The severity of obesity in AAV-Cre-injected mice had a good correlation with the extent of AAV infection in the PVH (Fig. 3d), but not along needle tracks or in other hypothalamic areas (Supplementary Table 1). These results show that PVH TrkB is critical for the control of energy balance.

**Figure 3.**
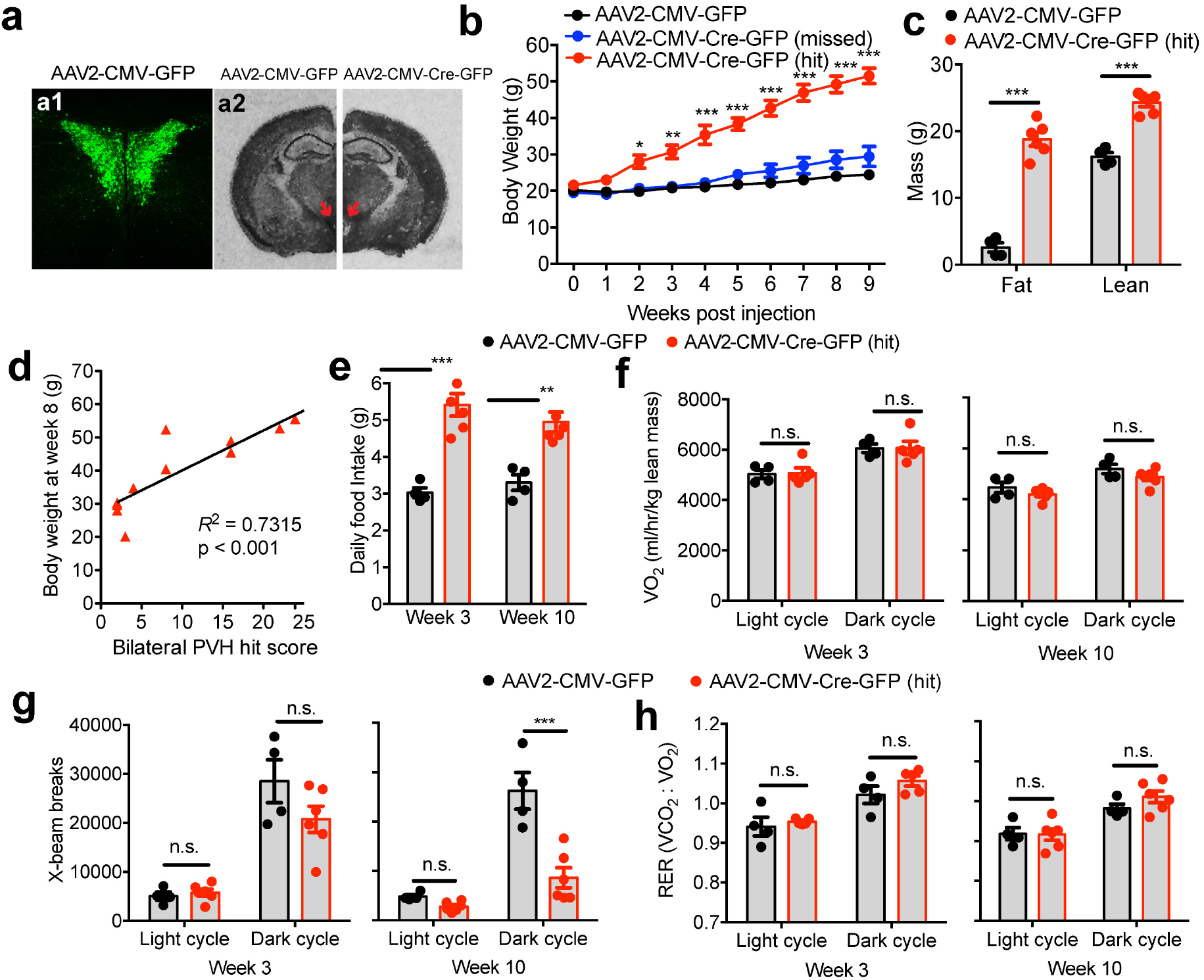
Deletion of the *Ntrk2* gene in the PVH of adult female mice leads to hyperphagic obesity. (**a**) Bilateral injection of AAV2-CMV-GFP or AAV2-CMV-Cre-GFP (200 nl) into the PVH of *Ntrk2*^*lox/lox*^ mice. GFP fluorescence indicates precise injection of AAV2-CMV-GFP into the PVH bilaterally (a1). In situ hybridization shows abolishment of *Ntrk2* expression in the PVH of *Ntrk2*^*lox/lox*^ mice injected with AAV2-CMV-Cre-GFP, but not AAV2-CMV-GFP (a2). Arrows denote the PVH. (**b**) Body weight of *Ntrk2*^*lox/lox*^ mice injected with either AAV2-CMV-GFP or AAV2-CMV-Cre-GFP into the PVH bilaterally. Mice injected with AAV2-CMV-Cre-GFP were divided into two groups: AAV2-CMV-Cre-GFP (missed) mice in which there was little viral transduction in the PVH and AAV2-CMV-Cre-GFP (hit) mice in which there was significant viral transduction in the PVH. n = 4 mice for AAV2-CMV-GFP, 5 mice for AAV2-CMV-Cre-GFP (missed), and 6 mice for AAV2-CMV-Cre-GFP (hit). Two-way ANOVA with post hoc Bonferroni multiple comparisons; *F*(2, 120) = 37.13, *P* < 0.0001 for viral injection; \*\**P* < 0.01 and \*\*\**P* < 0.001 for comparisons between AAV2-CMV-Cre-GFP (hit) and AAV2-CMV-GFP or AAV2-CMV-Cre-GFP (missed). (**c**) Body composition of *Ntrk2*^*lox/lox*^ mice 9 weeks after AAV injection. Student’s *t* test; \*\*\**P* < 0.001. (**d**) Correlation between the extent of AAV2-CMV-Cre-GFP transduction in PVH and body weight at 8 weeks after injection. Data are from both AAV2-CMV-Cre-GFP (hit) and AAV2-CMV-Cre-GFP (missed) groups. (**e**) Daily food intake of *Ntrk2*^*lox/lox*^ mice during week 3 and week 10 post injection. Two-way ANOVA with post hoc Bonferroni multiple comparisons; ** *p*<0.01 and *** *p*<0.001. (**f**) O_2_ consumption of *Ntrk2*^*lox/lox*^ mice during week 3 and week 10 post injection. Two-way ANOVA with post hoc Bonferroni multiple comparisons; n.s., no significant. (**g**) Locomotor activity of *Ntrk2*^*lox/lox*^ mice during week 3 and week 10 post injection. Two-way ANOVA with post hoc Bonferroni multiple comparisons; n.s. = not significant and \*\*\**P* < 0.001. (**h**) RER of *Ntrk2*^*lox/lox*^ mice during week 3 and week 10 post injection. Two-way ANOVA with post hoc Bonferroni multiple comparisons; n.s. = not significant. Error bars indicate SEM.

Mice in the AAV2-CMV-Cre-GFP (hit) group displayed marked hyperphagia just two weeks after AAV injection and remained hyperphagic 10 weeks after injection (Fig. 3e). These mice had energy expenditure and respiratory exchange ratio comparable to control mice (Fig. 3f, h), whereas their ambulatory activity was trending lower 2 weeks after injection and was reduced 10 weeks after injection (Fig. 3g). These results indicate that PVH TrkB regulates energy balance mainly by suppressing appetite.

We noticed that the extent of AAV infection along a needle track is correlated with injection volume. We sought to further verify the role of PVH TrkB in the control of appetite by injecting female *Ntrk2*^*lox/lox*^ mice with a smaller volume of AAV into the PVH to minimize infection in needle tracks and other hypothalamic nuclei. We detected very few AAV-transduced neurons in brain areas other than the PVH (mostly detected in Bregma −0.7 to −0.94 mm) in this batch of mice (Supplementary Fig. 3a1-a3). In the PVH, *Ntrk2* gene deletion did not cause neuronal loss during the course of the experiment (Supplementary Fig. 3a4-a6). Mice injected with AAV2-CMV-Cre-GFP displayed the same phenotypes as we observed in the first batch of mice, i.e. severe hyperphagic obesity, increased linear growth and reduced ambulatory activity (Supplementary Fig. 3b-g). Interestingly, even unilateral *Ntrk2* gene deletion was sufficient to cause significant obesity (Supplementary Fig. 3b), indicating the extreme importance of PVH TrkB in the control of energy balance. Furthermore, *Ntrk2* gene deletion in the PVH also led to obesity in male mice (Supplementary Fig. 3h). We did not detect any toxic effect on PVH neurons from AAV transduction, as bilateral injection of AAV2-CMV-Cre-GFP into the PVH of wild-type mice did not alter body weight (Supplementary Fig. 3i).

Oxytocin neurons in the PVH may play a central role in the control of appetite. Ablation of the oxytocin receptor-expressing neurons in the hindbrain leads to overeating^42^. Loss of oxytocin neurons is associated with both Prader-Willi syndrome and *SIM1* mutations, each of which leads to insatiable appetite and severe obesity in humans^43–46^. Because a large number of oxytocin neurons express TrkB in the PVH (Fig. 1b), we investigated whether TrkB signaling suppresses appetite through these neurons by deleting the *Ntrk2* gene in oxytocin neurons using the *Oxt*^*Cre*^ mouse strain^47^ (Supplementary Fig. 4a). *Ntrk2* deletion in oxytocin neurons did not affect body weight in mice of either sex (Supplementary Fig. 4b, c).

Collectively, these selective gene deletion experiments show that signaling through the TrkB receptor potently suppresses appetite through non-oxytocin-expressing PVH neurons. They also indicate that TrkB signaling in the PVH somewhat promotes locomotion without significantly altering energy expenditure.

### PVH^TrkB^ neurons bidirectionally regulate food intake

As BDNF potently potentiates neuronal growth and synaptic function^19^, *Ntrk2* deletion would impair the function of PVH^TrkB^ neurons. The observation that *Ntrk2* deletion in the PVH leads to hyperphagic obesity would implicate PVH^TrkB^ neurons in the acute regulation of food intake. We tested this hypothesis by manipulating the activity of PVH^TrkB^ neurons using designer receptors exclusively activated by designer drugs (DREADDs) and their ligand clozapine N-oxide (CNO)^48,49^. We expressed stimulatory DREADD hM3D(Gq), inhibitory DREADD hM4D(Gi), or control mCherry in PVH^TrkB^ neurons by injecting Cre-dependent AAV2-hSyn-DIO-hM3D(Gq)-mCherry, AAV2-hSyn-DIO-hM4D(Gi)-mCherry, or AAV2-hSyn-DIO-mCherry into the PVH of *Ntrk2*^*CreER*/+^ mice. We subsequently induced the expression of the injected AAV vectors in PVH^TrkB^ neurons by treating the mice with tamoxifen (Fig. 4a, b).

**Figure 4.**
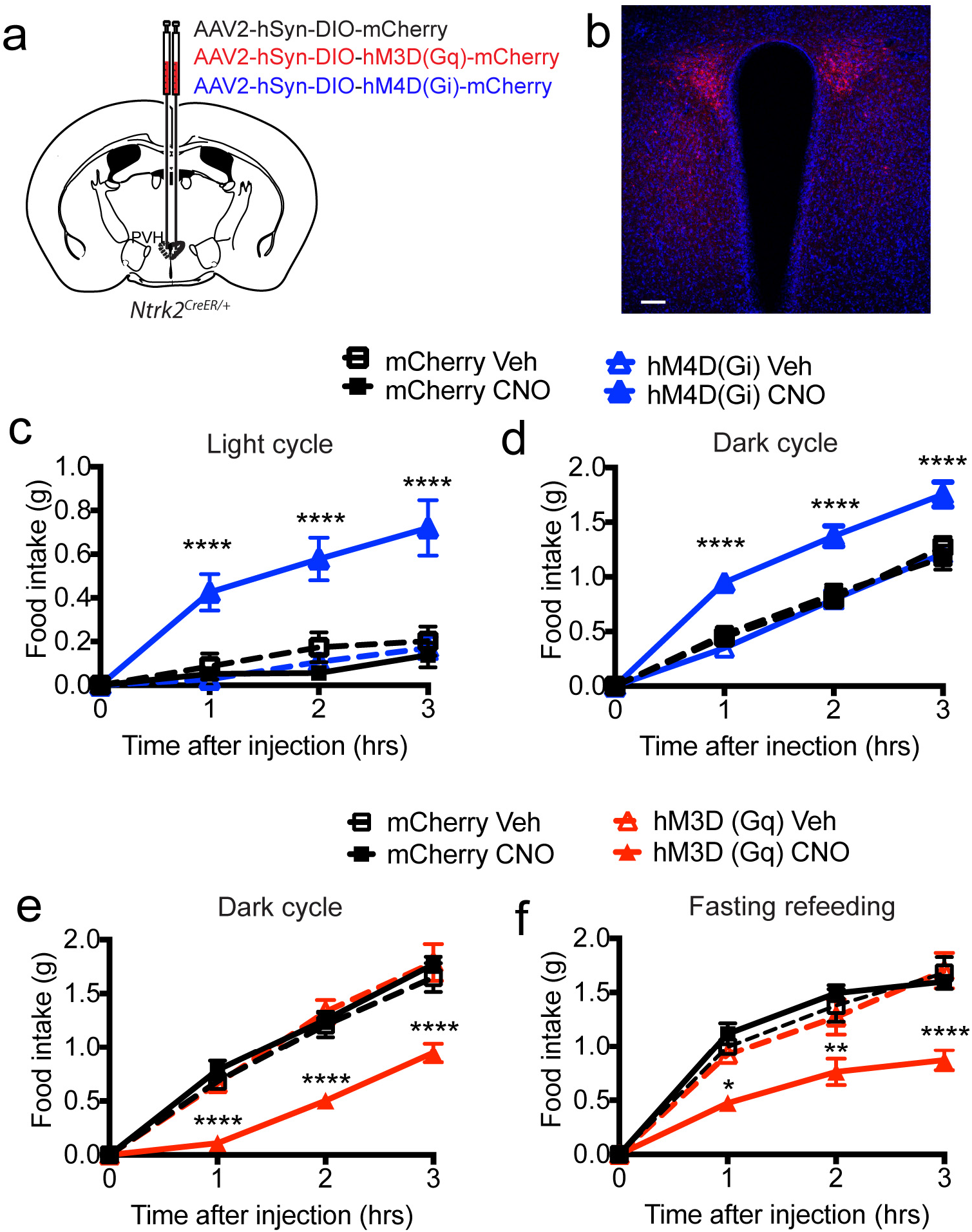
Chemogenetic modification of PVH^TrkB^ neuronal activity alters food intake. (**a**) A diagram showing bilateral injection of AAV2-hSyn-DIO-mCherry, AAV2-hSyn-DIO-hM3D(Gq)-mCherry, or AAV2-hSyn-DIO-hM4D(Gi)-mCherry into the PVH of *Ntrk2*^*CreER*/+^ mice. (**b**) A confocal imaging showing mCherry expression of injected AAV in the PVH. Brain sections were counter-stained with DAPI. The scale bar represents 100 μm. (**c**) Inhibition of PVH^TrkB^ neurons stimulated food intake during the light cycle. n = 5 and 6 mice for AAV2-hSyn-DIO-mCherry (mCherry) and AAV2-hSyn-DIO-hM4D(Gi)-mCherry [hM4D(Gi)] groups, respectively. Two-way ANOVA with post hoc Tukey’s multiple comparisons; *F*(3, 72) = 35.23, *P* < 0.0001 for virus and treatment; ****p < 0.0001 when compared to the hM4D(Gi)-vehicle (Veh) group. (**d**) Inhibition of PVH^TrkB^ neurons stimulated food intake during the dark cycle. n = 6 and 6 mice for mCherry and hM4D(Gi) groups, respectively. Two-way ANOVA with post hoc Tukey’s multiple comparisons; *F*(3, 76) = 31.49, *P* < 0.0001 for virus and treatment; ****p < 0.0001 when compared to the hM4D(Gi)-Veh group. (**e**) Activation of PVH^TrkB^ neurons suppressed food intake during the dark cycle. Two-way ANOVA with post hoc Bonferroni’s multiple comparisons; *F*(3, 22) = 17.27, *P* < 0.0001 for virus and treatment; \*\*\*\**P* < 0.0001 when compared to the hM3D(Gq)-Veh group. (**f**) Activation of PVH^TrkB^ neurons suppressed fasting-induced appetite. Two-way ANOVA with post hoc Bonferroni’s multiple comparisons; *F*(3, 22) = 10.47, *P* < 0.0001 for virus and treatment; \**P* < 0.05, \*\**P* < 0.01 and \*\*\*\**P* < 0.0001 when compared to the hM3D(Gq)-Veh group. Error bars indicate SEM.

Administration of CNO did not alter food intake of *Ntrk2*^*CreER*/+^ mice expressing mCherry during either the light or dark cycle or after fasting (Fig. 4c-f), indicating that CNO does not have any detectable nonspecific effect on feeding. Administration of CNO greatly increased food intake of *Ntrk2*^*CreER*/+^ mice expressing hM4D(Gi)-mCherry during the light cycle when mice do not normally eat much food, compared with vehicle administration (Fig. 4c). This result indicates that PVH^TrkB^ neurons are active to maintain physiological satiation during the light cycle. Interestingly, inhibition of PVH^TrkB^ neurons also increased food intake during the dark cycle when mice are physiologically hungry and ingest the vast majority of their daily energy intake (Fig. 4d), suggesting that the anorexigenic PVH^TrkB^ neurons are not fully silenced when mice are physiologically hungry. Conversely, CNO administration drastically reduced food intake of *Ntrk2*^*CreER*/+^ mice expressing hM3D(Gq)-mCherry during the dark cycle or after overnight fasting, compared with vehicle administration (Fig. 4e, f). This result indicates that activation of PVH^TrkB^ neurons is sufficient to suppress appetite. Taken together, these chemogenetic experiments indicate that PVH^TrkB^ neurons are partially silenced to allow for feeding when mice are hungry, while being active to maintain physiological satiation during the light cycle, and thus PVH^TrkB^ neurons play a critical role in the regulation of appetite by promoting satiety.

### PVH^TrkB^ neurons project to multiple brain areas

We sought to identify the downstream sites through which PVH^TrkB^ neurons regulate appetite. To map the projections of PVH^TrkB^ neurons in the brain, we injected Cre-dependent AAV2-CAG-FLEX-tdTomato into the PVH of adult *Ntrk2*^*CreER*/+^ mice (Fig. 5a1). After tamoxifen treatment, tdTomato nicely labeled PVH^TrkB^ neurons across the rostral-caudal axis (Fig. 5a2-4). PVH^TrkB^ neurons project to four main brain sites: the VMH (Fig. 5a5), the median eminence (ME; Fig. 5a5), the lateral parabrachial nucleus (LPBN; Fig. 5a6), and the nucleus tractus solitaris (NTS; Fig. 5a8). Lesser projections were detected in the medial parabrachial nucleus (MPBN; Fig. 5a6), the locus coeruleus (LC; Fig. 5a7), the dorsal motor nucleus of the vagus (DMV; Fig. 5a8), and the ventrolateral periaqueductal gray (vlPAG; Supplementary Fig. 5a). In the LPBN, axons of PVH^TrkB^ neurons mainly terminate at the central compartment, which is next to the CGRP (calcitonin gene-related peptide)-marked external compartment^50^ (Fig. 5b).

**Figure 5.**
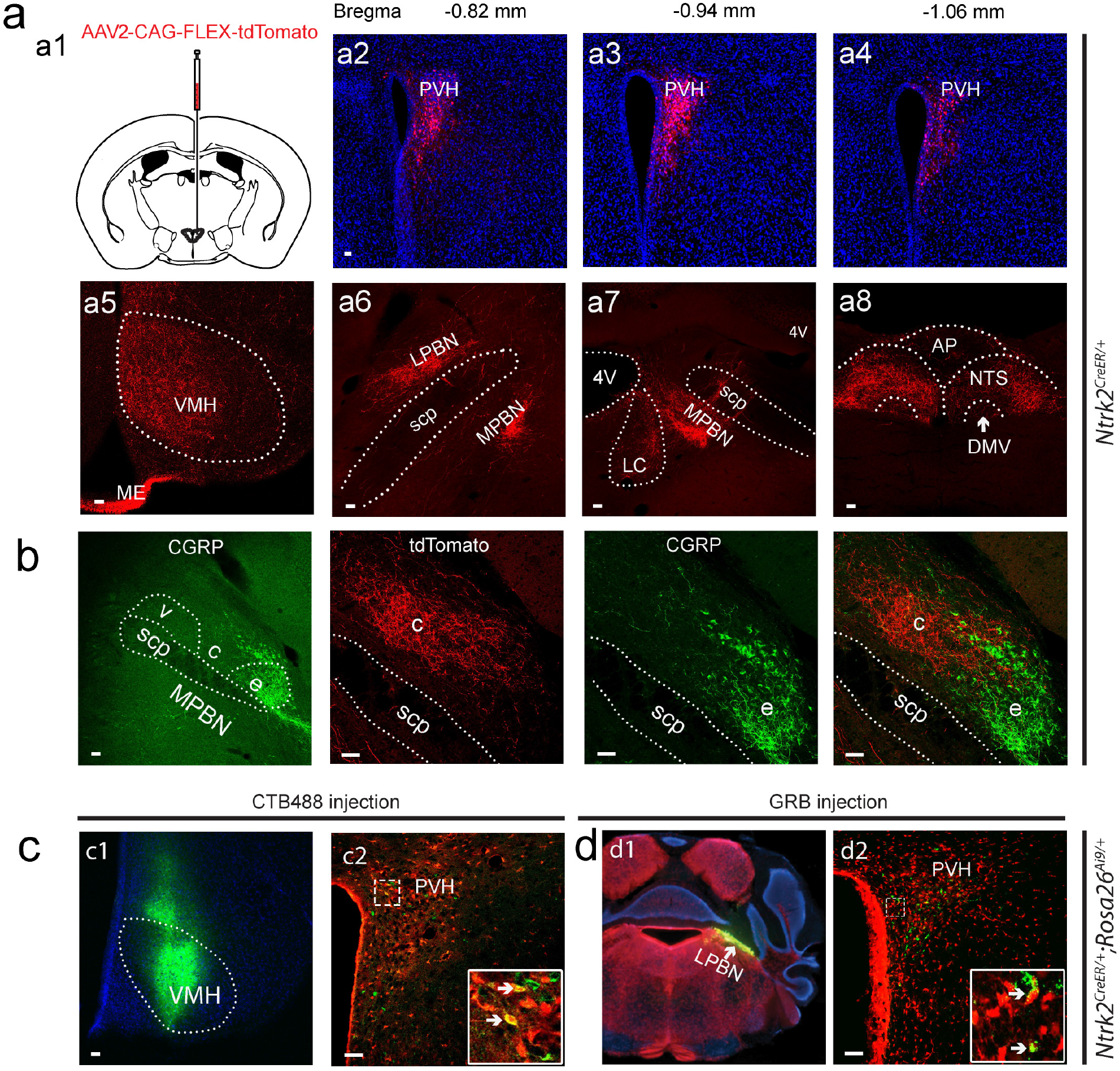
Projections of PVH^TrkB^ neurons. (**a**) AAV2-CAG-FLEX-tdTomato was unilaterally injected into the PVH of *Ntrk2*^*CreER*/+^ mice (a1). After the mice were treated with tamoxifen, tdTomato-labeled PVH^TrkB^ neurons were present throughout the rostral-caudal axis (a2-a4). Axonal terminals of these neurons were found in the VMH, LPBN, MPBN, LC, NTS and DMV (a5-a8). (**b**) The LPBN includes the external (e), central (c) and ventral (v) compartments. Axonal terminals of PVH^TrkB^ neurons are mainly in the central compartment, which is next to the CGRP-marked external compartment. (**c**& **d**) Retrograde tracers, CTB488 and GRB, were unilaterally injected into the VMH (c) or LPBN (d) of tamoxifen-treated *Ntrk2*^*CreER*/+^*;Rosa26*^*Ai9*/+^ mice and labeled some tdTomato-expressing PVH^TrkB^ neurons. AP, area postrema; DMV, dorsal motor nucleus of the vagus; ME, median eminence; LC, locus coeruleus; LPBN, lateral parabrachial nucleus; MPBN, medial parabrachial nucleus; NTS, nucleus tractus solitaris; PVH, paraventricular hypothalamus; scp, superior cerebellar peduncle; VMH, ventromedial hypothalamus. Scale bars represent 50 μm.

The projection to the ME is expected, as TrkB is expressed in a fraction of PVH neurons expressing GHRH (Supplementary Fig. 1d) and oxytocin (Fig. 1b), which project to the ME and the posterior pituitary via the ME, respectively, to release hormones. To validate other putative projection fields, we injected retrograde tracers, Alexa Fluor 488-conjugated cholera toxin subunit B (CTB488) or green retrobead (GRB), into each of these fields in tamoxifen-treated *Ntrk2*^*CreER*/+^*;Rosa26*^*Ai9*/+^ mice. In a true projection field, retrograde tracers would be taken up at axonal terminals and retrogradely transported to the cell bodies of PVH^TrkB^ neurons. Indeed, we found that retrograde tracers injected into the VMH (Fig. 5c), LPBN (Fig. 5d), vlPAG, LC, and NTS (Supplementary Fig. 5) labelled some TrkB-expressing neurons in the PVH. These results show that PVH^TrkB^ neurons project to the central compartment of the LPBN (cLPBN), VMH, NTS, MPBN, DMV, vlPAG, ME and pituitary.

### Projection-specific gene deletion

In order to determine at which projection PVH TrkB is necessary for the regulation of appetite, we designed a two-virus system for projection-specific gene deletion. In this strategy, one viral vector is a retrograde virus [canine adenovirus 2 (CAV2)^51^ or retrograde AAV (AAVretro)^52^] expressing yeast flippase (FLP), whereas the other viral vector is AAV2-Ef1a-fDIO-mCherry-P2A-Cre, in which expression of mCherry-P2A-Cre is dependent on FLP-mediated recombination at two sets of Frt sequences (Fig. 6a). If we stereotaxically inject FLP-expressing retrograde virus into the LPBN and AAV2-Ef1a-fDIO-mCherry-P2A-Cre into the PVH, the retrograde virus would be taken up at axonal terminals and retrogradely transported to the cell bodies of neurons that project to the LPBN. In the PVH, FLP expressed in neurons that project to the LPBN would mediate recombination between the two sets of Frt sites in AAV2-Ef1a-fDIO-mCherry-P2A-Cre and invert the orientation of the mCherry-P2A-Cre sequence, leading to the expression of mCherry and Cre (Fig. 6a). If the two viral vectors are injected into *Ntrk2*^*lox/lox*^ mice, expressed Cre would delete the *Ntrk2* gene in PVH neurons that project to the LPBN.

**Figure 6.**
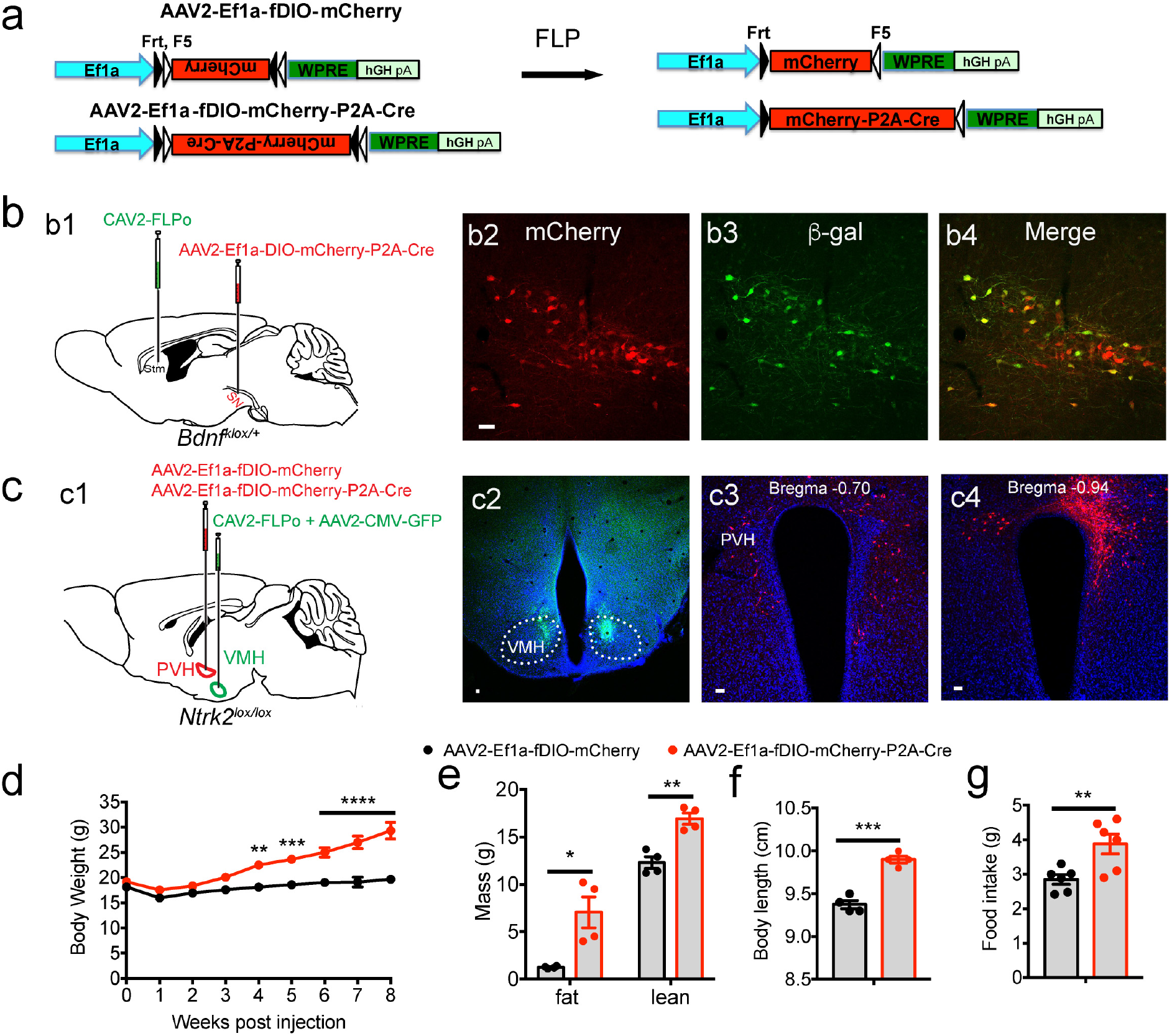
Deletion of the *Ntrk2* gene in PVH neurons projecting to the VMH leads to hyperphagia and obesity. (**a**) Diagrams of AAV2-Ef1a-fDIO-mCherry and AAV2-Ef1a-fDIO-mCherry-P2A-Cre viral vectors for FLP-dependent expression of mCherry and Cre. (**b**) Deletion of the *Bdnf* gene in substantia nigra neurons projecting to the striatum. CAV2-FLPo (500 nl) and AAV2-Ef1a-fDIO-mCherry-P2A-Cre (200 nl) were stereotaxically injected into the striatum (Stm) and substantia nigra (SN) of *Bdnf*^*klox*/+^ mice, respectively (b1). Expression of mCherry and β-galactosidase (β-gal) was detected in the substantia nigra. (**c**) Deletion of the *Ntrk2* gene in PVH neurons projecting to the VMH. A mixture of CAV2-FLPo + AAV2-CMV-GFP (350 nl, 5:1 ratio) and either AAV2-Ef1a-fDIO-mCherry (control) or AAV2-Ef1a-fDIO-mCherry-P2A-Cre (150 nl) were bilaterally injected into the VMH and PVH of *Ntrk2*^*lox/lox*^ mice, respectively (c1). Co-injected AAV2-CMV-GFP was used to mark CAV2 injection sites (c2). Neurons that are transduced by both CAV2-FLPo and one of the two FLP-dependent AAV vectors would express mCherry (c3 and c4). (**d-g**) Body weight (d), fat mass and lean mass (e), body length (f), and food intake (g) of female *Ntrk2*^*lox/lox*^ mice injected with CAV2-FLPo into the VMH and either AAV2-Ef1a-fDIO-mCherry or AAV2-Ef1a-fDIO-mCherry-P2A-Cre into the PVH. Body weight data were analyzed using two-way ANOVA with Bonferroni *post hoc* tests; *F*_(1, 90)_ = 124.0, *P* < 0.0001 for virus. Other data were analyzed with Student’s *t* test. n = 6 mice for each group. \**P* < 0.05, \*\**P* < 0.01, \*\*\**P* < 0.001 and \*\*\*\**P* < 0.0001. Scale bars represent 50 μm.

We conducted a proof-of-principle experiment to test this projection-specific gene deletion approach in the well-defined substantia nigra → striatum projection. We injected CAV2-FLPo, which expresses optimized FLP, into the striatum and AAV2-Ef1a-fDIO-mCherry-P2A-Cre into the substantia nigra of *Bdnf*^*klox*/+^ mice (Fig. 6b1). *Bdnf*^*klox*/+^ mice contains a uniquely floxed *Bdnf* allele that starts to express β-galactosidase in BDNF-expressing cells when Cre recombines the allele^53^. We detected many mCherry-expressing neurons in the substantia nigra (Fig. 6b2), indicating that CAV2-FLPo was retrogradely transported to the substantia nigra from the striatum and induced mCherry expression from AAV2-Ef1a-fDIO-mCherry-P2A-Cre. Because mCherry and Cre are in the same cistron, mCherry-positive neurons should also express Cre, which would mediate recombination of the *Bdnf*^*klox*^ allele. Indeed, we found that some mCherry-positive neurons expressed β-galactosidase in the substantia nigra (Fig. 6b2-4), where ~50% dopaminergic neurons express BDNF^54^. As expected, we detected expression of neither mCherry nor β-galactosidase in the substantia nigra when CAV2-FLPo was not injected into the striatum (Supplementary Fig. 6a-d). This experiment shows that our two-virus system works for projection-specific gene deletion.

### Deletion of the *Ntrk2* gene in PVH neurons projecting to the VMH and LPBN leads to hyperphagia and obesity

We targeted the *Ntrk2* gene in PVH neurons that project to the NTS, VMH and LPBN, the three main projection fields of PVH^TrkB^ neurons. Retrograde virus CAV2-FLPe-GFP (expressing a fusion protein of GFP and the improved FLP), CAV2-FLPo, or AAV2retro-hSyn-FLPo-T2A-GFP were used to express FLP in cell bodies of afferent neurons. We conducted this study only in female mice. Because *Ntrk2* deletion in the PVH produced more severe obesity in females than males (Fig. 3b and Supplementary Fig. 3h), we used female mice to more sensitively uncover the role of different PVH^TrkB^ projections in the control of appetite.

To delete the *Ntrk2* gene in PVH neurons projecting to the NTS (PVH^TrkB→NTS^ neurons), we injected CAV2-FLPe-GFP into the NTS and either AAV2-Ef1a-fDIO-mCherry (control) or AAV2-Ef1a-fDIO-mCherry-P2A-Cre into the PVH in *Ntrk2*^*lox/lox*^ mice (Supplementary Fig. 6e). Because the NTS is a long structure, we infused a mixture of CAV2-FLPe-GFP and GRBs into two sites, the rostral part (Supplementary Fig. 6f1-3) and the caudal part (Supplementary Fig. 6f4-6) of the central NTS. Co-injected GRBs allows us to determine if an injection hits the NTS (Supplementary Fig. 6f). In hit mice, we detected mCherry expression in the PVH (Supplementary Fig. 6g), indicating expression of Cre in AAV2-Ef1a-fDIO-mCherry-P2A-Cre injected mice, which would delete the *Ntrk2* gene in positive neurons. We found comparable body weight (Supplementary Fig. 6h), body composition (Supplementary Fig. 6i) and food intake (Supplementary Fig. 6j) between mutant mice and control mice. These results suggest that TrkB in PVH^TrkB→NTS^ neurons is not essential for the control of food intake and body weight.

We next targeted the *Ntrk2* gene in PVH neurons projecting to the VMH (PVH^TrkB→VMH^ neurons) by injecting a mixture of CAV2-FLPo and AAV-GFP into the VMH and either AAV2-Ef1a-fDIO-mCherry or AAV2-Ef1a-fDIO-mCherry-P2A-Cre into the PVH in *Ntrk2*^*lox/lox*^ mice (Fig. 6c1). Co-injected AAV-GFP marks injection sites (Fig. 6c2). In our initial trial, we noticed lesions at CAV2-FLPo injection sites, likely due to an immune response to CAV2. Therefore, we treated *Ntrk2*^lox/lox^ mice with the immunosuppressant cyclophosphamide (50 mg/kg, i.p.) once every 2 d from day −3 to day 13 as previously described^55^. Ten weeks after CAV2 injection, we did not detect any lesions in the VMH (Fig. 6c2), indicating that cyclophosphamide successfully blocked the immune response to CAV2. In mice with successful viral injection as evidenced by bilateral mCherry expression in the PVH (Fig. 6c3 and 6c4), the Cre group had significantly higher body weight (Fig. 6d), fat mass and lean mass (Fig. 6c), body length (Fig. 6f), and food intake (Fig. 6g). These results show that deleting *Ntrk2* in PVH^TrkB→VMH^ neurons leads to hyperphagia and obesity, indicating a crucial role for TrkB in PVH^TrkB→VMH^ neurons in appetite suppression.

Finally, we deleted the *Ntrk2* gene in PVH neurons projecting to the LPBN (PVH^TrkB→LPBN^ neurons) by injecting AAV2retro-hSyn-FLPo-T2A-GFP into the LPBN and either AAV2-Ef1a-fDIO-mCherry or AAV2-Ef1a-fDIO-mCherry-P2A-Cre into the PVH in *Ntrk2*^*lox/lox*^ mice (Fig. 7a1). We employed AAV2retro-hSyn-FLPo-T2A-GFP rather than CAV2-FLPo for the PVH → LPBN projection, because we found that AAVretro is more efficient in retrograde infection than CAV2 at this projection. AAV2retro-hSyn-FLPo-T2A-GFP expresses GFP at injection sites (Fig. 7a2), allowing us to assess the accuracy of each injection. In mice with successful viral injection as evidenced by GFP expression in the LPBN (Fig. 7a2) and mCherry expression in the PVH (Fig. 7a3 and 7a4), the Cre group had significantly higher body weight (Fig. 7b), fat mass and lean mass (Fig. 7c), body length (Fig. 7d), and food intake (Fig. 7e). These results show that deleting *Ntrk2* in PVH^TrkB→LPBN^ neurons also leads to hyperphagia and obesity, indicating a crucial role for TrkB in PVH^TrkB→LPBN^ neurons in the control of food intake.

**Figure 7.**
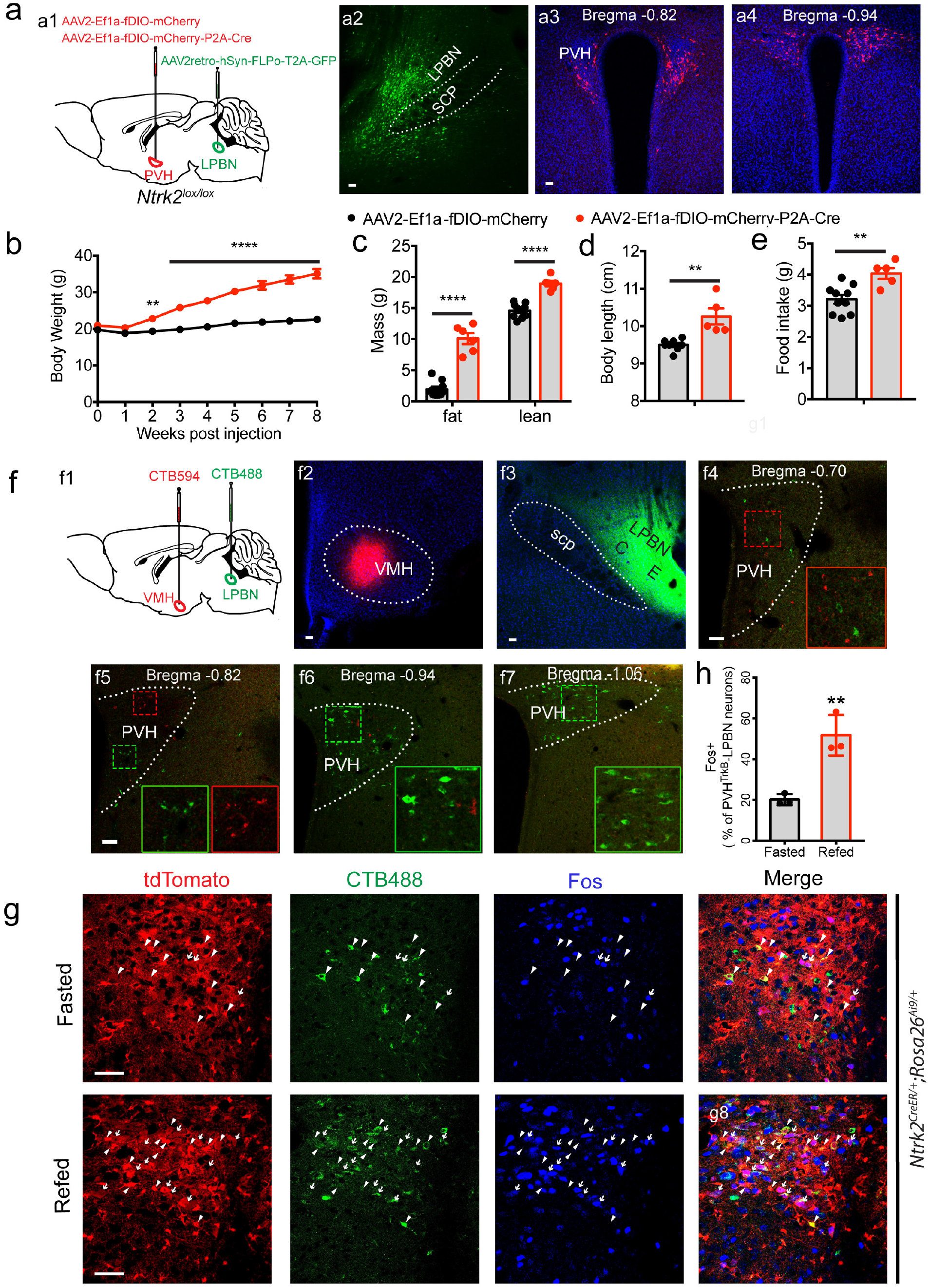
PVH neurons projecting to the LPBN are activated by refeeding and their TrkB expression is crucial for appetite suppression. (**a**) Deletion of the *Ntrk2* gene in PVH neurons projecting to the LPBN. AAV2retro-hSyn-FLPo-T2A-GFP (200 nl) and either AAV2-Ef1a-fDIO-mCherry (control) or AAV2-Ef1a-fDIO-mCherry-P2A-Cre (150 nl) were bilaterally injected into the LPBN and PVH of *Ntrk2*^*lox/lox*^ mice, respectively (a1). GFP marks injection sites of AAV2retro-hSyn-FLPo-T2A-GFP (a2). Neurons that are transduced by both AAV2retro-hSyn-FLPo-T2A-GFP and one of the two FLP-dependent AAV vectors would express mCherry (a3 and a4). (**b**-**e**) Body weight (b), fat mass and lean mass (c), body length (d), and food intake (e) of female *Ntrk2*^*lox/lox*^ mice injected with AAV2retro-hSyn-FLPo-T2A-GFP into the LPBN and either AAV2-Ef1a-fDIO-mCherry or AAV2-Ef1a-fDIO-mCherry-P2A-Cre into the PVH. Body weight data were analyzed using two-way ANOVA with Bonferroni *post hoc* tests; *F*(1, 126) = 488.7, *P* < 0.0001 for virus. Other data were analyzed with Student’s *t* test. n = 10 mice in the control group and 6 mice in the Cre group. \*\**P* < 0.01 and \*\*\*\**P* < 0.0001. (**f**) Retrogradely labeling of PVH neurons. CTB594 and CTB488 were injected into the VMH and the LPBN, respectively (f1-3). Some retrogradely labeled neurons are in the PVH (f4-7). (**g & h**) Refeeding after overnight food deprivation induced Fos expression in PVH^TrkB^ neurons projecting to the LPBN. Confocal images show colocalization of tdTomato, CTB488 and Fos in the PVH of tamoxifen-treated *Ntrk2*^*CreER*/+^;*Rosa26*^*Ai9*/+^ mice (g). In these images, tdTomato marks TrkB neurons, CTB488 labels neurons projecting to the LPBN, and Fos indicates activated neurons. Arrowheads denote neurons positive for tdTomato and CTB488, while arrows indicate neurons positive for all three markers. Number of Fos^+^ neurons in PVH^TrkB→LPBN^ neurons under fasting and refeeding conditions was quantified (h). Scale bars represent 50 μm.

### PVH^TrkB^ neurons projecting to the LPBN are distinct from those projecting to the VMH and respond to nutritional state

Because the TrkB receptor at both PVH^TrkB^ projections to the VMH and the LPBN is crucial for the control of appetite, we next investigated the possibility that the same subset of PVH^TrkB^ neurons send collaterals to the VMH and the LPBN. We stereotaxically injected retrograde tracers CTB488 and Alexa Fluor 594-conjugated CTB (CTB594) into the VMH and LPBN, respectively (Fig. 7f1-3). We found that CTB594 mainly labeled cell bodies in the rostral and medial parts of the PVH, whereas CTB488 largely in the medial and caudal parts of the PVH (Fig. 7f4-7). No neurons were detected to contain both CTB488 and CTB594 (Fig. 7f4-7). These results indicate that two distinct subtypes of PVH^TrkB^ neurons project to the VMH and the LPBN.

The observation that abolishment of *Ntrk2* expression in PVH^TrkB→VMH^ and PVH^TrkB→LPBN^ neurons leads to hyperphagic obesity suggests that these two subtypes of neurons respond to changes in nutritional state. We tested this hypothesis in PVH^TrkB→LPBN^ neurons. We injected CTB488 into the LPBN of tamoxifen-treated *Ntrk2*^*CreER*/+^;*Rosa26*^*Ai9*/+^ mice. After the mice were extensively handled, they were fasted overnight and a half of them were refed for 2 h. We found that more CTB488-labeled PVH^TrkB^ neurons expressed Fos in refed mice than in fasted mice (Fig. 7g, h), indicating that refeeding activates PVH^TrkB→LPBN^ neurons. These results together with the findings from *Ntrk2* deletion and manipulation of neuronal activity, shows that feeding activates PVH^TrkB→VMH^ and PVH^TrkB→LPBN^ neurons to suppress appetite.

## DISCUSSION

This study has identified the PVH as a main site where TrkB signaling works to suppress food intake. Using a projection-specific gene deletion strategy we designed, we have further narrowed down the action site of TrkB signaling to two distinct subtypes of PVH neurons that project to the LPBN and the VMH, respectively. Given that BDNF-TrkB signaling is known to potentiate synaptic transmission in the hippocampus^18,19^, these results suggest that some PVH^TrkB^ neurons are activated to suppress appetite. Indeed, we have found in mice that refeeding activates PVH^TrkB→LPBN^ neurons and that chemogenetic activation of PVH^TrkB^ neurons inhibits food intake during the dark cycle or after overnight food deprivation. Thus, our study indicates that signals reflecting repletion activate PVH^TrkB^ neurons to promote satiety through their outputs to the LPBN and the VMH.

We found that deletion of the *Ntrk2* gene with the Sim1-Cre transgene led to hyperphagia and reduced energy expenditure in mice. This observation is in agreement with the previous finding that BDNF expressed in the PVH suppresses food intake and promotes adaptive thermogenesis^22^. Collectively, these results demonstrate that BDNF-TrkB signaling plays a critical role in the control of both energy intake and energy expenditure. When we selectively deleted the *Ntrk2* gene in the PVH using stereotaxic injection of Cre-expressing AAV, mutant mice displayed marked hyperphagia but normal energy expenditure. It is likely that TrkB signaling in other brain areas that express Sim1-Cre is involved in the regulation of energy expenditure. One of these brain areas could be the DMH, in which *Ntrk2* deletion significantly reduces energy expenditure^28^. Injection of BDNF protein into the PVH was reported to increase energy expenditure^37^. This could be a result of activation of the TrkB receptor at axonal terminals of neurons that project to the PVH or a non-specific effect of recombinant BDNF.

A genetically defined group of neurons in a specific brain structure usually have multiple projection fields, and the powerful optogenetic approach is commonly employed to uncover the role of the neurons at each projection in animal behaviors. For example, Garfield et al. expressed channelrhodopsin-2 (ChR2) in PVH^MC4R^ neurons and found that light activation of ChR2 at PVH^MC4R^ axonal terminals in the LPBN inhibited food intake, indicating that the PVH^MC4R^ → LPBN neurocircuit promotes satiety^27^. However, this approach is not suitable for uncovering gene function and is not ideal for examination of the long-term effect of altering neuronal activity. The projection-specific gene deletion described here makes it possible to determine the function of a gene in a specific neurocircuit. If deletion of a gene also alters the function of the targeted neurocircuit, the same experiment will also reveal both short-term and long-term impacts of altering the activity of the circuit on behaviors and physiology.

Using the projection-specific gene deletion approach, we have identified two subtypes of PVH^TrkB^ neurons (PVH^TrkB→LPBN^ neurons and PVH^TrkB→VMH^ neurons) in which TrkB signaling is necessary to suppress appetite. Deletion of the *Ntrk2* gene in one of these two subtypes of neurons led to hyperphagia and obesity. Furthermore, we have found that refeeding activates PVH^TrkB→LPBN^ neurons and that chemogenetic stimulation/inhibition of PVH^TrkB^ neurons greatly decreases/increases food intake, respectively. These results suggest that abolishment of TrkB signaling impairs the function of these two subtypes of neurons, which leads to deficits in transmitting satiation signals to their target neurons in the LPBN and VMH. These results also indicate that the PVH^TrkB^→LPBN and PVH^TrkB^→VMH neurocircuits are satiating and that BDNF is a key modulator of the two anorexigenic neurocircuits. As BDNF potently potentiates synaptic transmission^19^, it is likely that TrkB signaling enhances synaptic transmission in the two subtypes of PVH^TrkB^ neurons. It would be important to understand which neurons activate PVH^TrkB^ neurons in response to refeeding, what are the targets of PVH^TrkB^ neurons in the LPBN and VMH, and how TrkB signaling modulates synaptic function in PVH^TrkB→LPBN^ and PVH^TrkB→VMH^ neurons in future studies.

The vast majority of PVH neurons are glutamatergic neurons^56^. These neurons have been shown to provide excitatory drive to the AgRP neurons in the arcuate nucleus^57^, the cLPBN^27^, and the pre-locus coeruleus (pLC)^28^ to regulate satiety. This study identifies one more neurocircuit out of the PVH, i.e. PVH^TrkB^→VMH, to regulate satiety.

Although ablation of the oxytocin receptor-expressing neurons in the hindbrain leads to overeating^42^, ablation of PVH oxytocin neurons in adult mice has no effect on body weight, food intake, or energy expenditure^47^. Consistent with this observation, we found that deletion of the *Ntrk2* gene in PVH^OXT^ neurons did not affect energy balance. Thus, PVH^TrkB→LPBN^ and PVH^TrkB→VMH^ neurons should be distinct from oxytocin neurons. Previous work shows that LPBN-projecting PVH^MC4R^ (PVH^MC4R→LPBN^) neurons and PVH^BDNF^ neurons also promote satiety^22,27^, although it remains unknown where PVH^BDNF^ neurons project to suppress appetite. As we detected low co-localization of TrkB with either MC4R or BDNF, PVH^TrkB→LPBN^ and PVH^TrkB→VMH^ neurons could be distinct from PVH^MC4R→LPBN^ and PVH^BDNF^ neurons. A recent study shows that PDYN-expressing pLC-projecting PVH (PVH^PDYN→pLC^) neurons regulate satiety and are distinct from PVH^MC4R^ neurons^28^. If PVH^BDNF^ neurons are distinct from PVH^PDYN→pLC^ neurons, at least 5 subtypes of PVH neurons (PVH^MC4R→LPBN^, PVH^BDNF^, PVH^PDYN→pLC^, PVH^TrkB→LPBN^, and PVH^TrkB→VMH^) have been identified to promote satiety. It would be important to investigate how these neurons coordinately regulate satiety and body weight. This is especially true for PVH^MC4R→LPBN^ and PVH^TrkB→LPBN^ neurons, which project to the same target field, the cLPBN. It would be intriguing to know whether these two subtypes of neurons innervate the same neurons in the cLPBN and receive inputs from the same afferent neurons. These future studies will provide insights into the organization and function of the neural network that regulates appetite.

## ACKNOWLEDGEMENTS

We thank David Ginty for the *Ntrk2*^*CreER*^ mouse strain, Martin Wessendorf for the anti-TRH antibody, and Jessica Houtz and Shaw-wen Wu for critical reading of this manuscript. This work was supported by US National Institutes of Health grants to BX (R01 DK103335 and R01 DK105954).

## AUTHOR CONTRIBUTIONS

J.J.A. and B.X. conceived the study. J.J.A., C.E.K. and B.X. designed the study. J.J.A. did colocalization immunohistochemistry and analyzed Sim1-Cre;*Ntrk2*^*lox/lox*^ conditional knockout mice. C.E.K., J.J.A. and G.Y.L. did stereotaxic injection of AAV-Cre-GFP. C.E.K. and J.J.A. analyzed PVH-specific *Ntrk2* mutant mice. J.J.A. constructed viral vectors for projection-specific gene deletion, did DREADD experiments, and examined Fos induction after refeeding. J.J.A. constructed pCAV2-FLPe-GFP. E.J.K. produced CAV2-FLPe-GFP and CAV2-FLPo. J.J.A. and C.E.K. did projection tracing and projection-specific *Ntrk2* deletion. J.J.A. and B.X. prepared figures. B.X. and J.J.A. wrote the manuscript with input from all authors.

## METHODS

### Animals

Floxed *Ntrk2* (*Ntrk2*^*lox*^; also known as TrkB^lox^)^41^, floxed *Ntrk2*-*LacZ* (*Ntrk2*^*fBZ*^; also known as fBZ)^40^, *and* floxed *Bdnf* (*Bdnf*^*klox*^)^53^ mouse strains were previously described. *Ntrk2*^*lox*^ mice were backcrossed to C57BL/6J mice for at least 15 generations before they were used in this study. Sim1-Cre (stock No: 006395), MC4R-tau-GFP (stock No: 008323), *Rosa26*^*Ai9*^ [*Gt9(ROSA)26Sor*^*tm99CAG-tdTomato)Hze*^/J; stock No: 007909] and C57BL6/J (stock No: 000664) mouse strains were obtained from the Jackson Laboratory. The *Ntrk2*^*CreER*/+^ (also known as TrkB^CreER^) mouse strain^32^ was kindly provided by Dr. David Ginty at Harvard Medical School. We generated *Ntrk2*^*CreER*/+^*;Rosa26*^*Ai9*/+^ mice by crossing *Ntrk2*^*CreER*/+^ mice to *Rosa26*^*Ai9*/+^ mice. To induce nuclear localization of the CreER fusion protein in mice harboring the *Ntrk2*^*CreER*^ allele, we treated the mice with tamoxifen (2 mg/animal, i.p.) once a day for 7 consecutive days in 6-8-wk-old mice. Animals were used for experiments 7 d after the last tamoxifen injection. *Ntrk2*^*lox/lox*^ mice were crossed with Sim1-Cre*;Ntrk2*^*lox/lox*^ mice to generate Sim1-Cre*;Ntrk2*^*lox/lox*^ mutant mice and *Ntrk2*^*lox/lox*^ control mice. All mice used in this study were maintained in the C56BL6/J genetic background. Mice were maintained on a 12-h/12-h light/dark cycle with *ad libitum* access to water and regular laboratory chow (Teklad Rodent Diet 2019; 3.3 kcal/g energy density, 22% kcal from fat) unless otherwise specified. The Animal Care and Use Committees at The Scripps Research Institute Florida approved all animal procedures used in this study.

### Immunohistochemistry

For neuropeptide immunohistochemistry, we injected *Ntrk2*^*CreER*/+^*;Rosa26*^*Ai9*/+^ mice with colchicine (20 μg per animal, i.c.v.; Tocris) to impair protein transportation in neuronal processes. Forty-eight hours after colchicine injection, mice were deeply anaesthetized with avertin and transcardially perfused with phosphate buffered saline (PBS), followed by 4% paraformaldehyde in PBS. Coronal brain sections (40 μm in thickness) were obtained using a sliding microtome (Leica SM2000R). Brain sections were rinsed once with Tris buffered saline (TBS; 10 mM Tris-HCl, 150 mM NaCl, pH7.5), incubated with blocking buffer (0.4% Triton X-100, 1% bovine serum albumin, and 10% goat serum or horse serum in TBS) for 1 h at room temperature, and then incubated with primary antibodies diluted in the blocking buffer overnight at room temperature. The following primary antibodies were used: chicken anti-β-galactosidase (1:3,000; abcam #ab9361), rabbit anti-NeuN (1:1000; Millipore #ABN78), rabbit anti-GFAP (1:400; Sigma #G9269), mouse anti-oxytocin (1:5000; Millipore #MAB5296), rabbit anti-CRH (1:500; Millipore #AB1760), mouse anti-GFP (1:1000; Clontech #632460), rabbit anti-somatostatin (1:500; immunostar #20067), rabbit anti-GHRH (1:500; Immunostar #22938), rabbit anti-vasopressin (1:5000; Millipore #AB1565), mouse anti-tyrosine hydroxylase (1:10,000; Sigma #T1299), rabbit anti-c-Fos antibody (1:5000; abcam #ab208942), rabbit anti-prodynorphin (1:200; abcam #ab11137), and rabbit anti-TRH (1:10,000; generous gift from Dr. Martin Wessendorf, University of Minnesota). After three washes in TBS, sections were incubated with appropriate fluorescent secondary antibodies (1;500; Jackson ImmunoResearch) for 1 h at room temperature. Sections were washed three times in TBS, mounted onto slides, and coverslipped with a mounting medium containing 4’,6-diamidino-2-phenylindole (DAPI; Vector Laboratories). Fluorescent images were captured using a Nikon C2+ confocal microscope or a Leica TCS SP8 microscope.

### Plasmids and viruses

For generation of pAAV-Ef1a-fDIO-mCherry and pAAV-Ef1a-fDIO-mCherry-P2A-Cre, pAAV-Ef1a-fDIO-EYFP (a gift from Karl Deisseroth; Addgene plasmid #55641) was used as backbone. pAAV-Ef1a-fDIO-EYFP was digested with AscI and NheI site to remove EYFP. The sequence encoding mCherry and mCherry-P2A-Cre were obtained by PCR using plasmid pLM-CMV-R-Cre (a gift from Michel Sadelain; Addgene plasmid #27546) as template. PCR products were digested with AscI and NheI and cloned into the backbone of pAAV-Ef1a-fDIO-EYFP to generate plasmid pAAV-Ef1a-fDIO-mCherry and pAAV-Ef1a-fDIO-mCherry-P2A-Cre. pAAV-hSyn-FLPo-T2A-GFP was generated using pAAV-hSynapsin-Flpo (a gift from Massimo Scanziani; Addgene plasmid #60663) as backbone. We replaced the FLPo sequence containing a stop codon with a FLPo without a stop codon using BamHI and HindIII sites to generate an intermediate plasmid. The T2A-GFP sequence was amplified by PCR using plasmid PX461 (a gift from Feng Zhang; Addgene plasmid #48140) as template and inserted into the intermediate plasmid at HindIII site to generate pAAV-hSyn-FLPo-T2A-GFP. AAV2-Ef1a-fDIO-mCherry, AAV2-Ef1a-fDIO-mCherry-P2A-Cre and AAV2retro-hSyn-Flpo-T2A-GFP were packaged by UNC Vector Core. AAV2-CMV-GFP, AAV2-CMV-GFP-Cre and AAV2-CAG-FLEX-tdTomato viruses were purchased from UNC vector Core. AAV2-hSyn-DIO-mCherry, AAV2-hSyn-DIO-hM3Dq-mCherry and AAV2-hSyn-DIO-hM4Di-mCherry were purchased from Addgene.

To construct pCAV2-FLPe-GFP, we inserted the Flpe-GFP sequence into the CAV shuttle plasmid^58^ to generate pShuttle-FLPe-GFP. The FLPe-GFP sequence was isolated from pCAG-Flpe-GFP (a gift from Connie Cepko, Addgene plasmid #13788). The AscI-PacI fragment of the shuttle construct was purified and used to generate pCAV2-FLPe-GFP through homologous recombination in E. coli as described^59^. To generate pCAV2-FLPo, we began with pCAV-FLEx^loxP^-Flp^60^. pCAV-FLEx^loxP^-Flp contains the CAV-2 genome deleted in the E1 and E3 regions, and a cassette containing a CMV promoter, two loxP sites, an inverted FLPo open reading frame, and an SV40 polyA signal. pCAV-FLEx^loxP^-Flp was incubated with recombinant Cre recombinase (New England Biolabs) and then transformed into DHα bacteria. A clone containing the FLPo cassette in the 5’ → 3’ orientation was identified by restriction digestion analyses and denoted pCAV2-FLPo. Viral vectors CAV2-FLPe-GFP and CAV2-FLPo were generated, amplified, and purified as previously described^58^.

### Stereotaxic injection of virus and CTB

The stereotaxic surgeries were performed as previously described^22^. Mice were deeply anesthetized with isoflurane and were placed onto a stereotaxic apparatus (David Kopf, Model 940). A small incision was made to expose the skull, and a small hole was drilled on skull above the injection site. A Nanofil 33-gauge needle (World Precision Instruments, #NF33BV-2) was inserted into the brain and the virus was injected at a rate of 25 nl/min using Microsyringe Pump (World Precision Instruments, Micro4). After infusion, the needle was stayed in the same position for 5 min. Following injections, mice received subcutaneous injection of Loxicom (0.5 mg/kg) to relieve pain. Stereotaxic coordinates from the bregma used for this study were PVH (AP: −0.5 mm, ML: ±0.25 mm, DV: −5.4 mm), VMH (AP: −1.4 mm, ML: ±0.4 mm, DV: −5.95 mm), LPBN (AP: −5 mm, ML: ±1.5 mm, DV: −3.8 mm), vlPAG (AP: −4.72 mm, ML: ±0.68 mm, DV: −2.70 mm), and LC (AP: −5.4 mm, ML: ±0.91 mm, DV: −4.1 mm). Stereotaxic coordinates for the NTS relative to the calamus scriptorius were rostral NTS (AP: +0.85 mm, ML: ±0.58 mm, DV: −0.42 mm) and caudal NTS (AP: −0.0 mm, ML: ±0.26 mm, DV: −0.4 mm). After physiological measurements are completed, animals were perfused with saline and 4% paraformaldehyde and their brains were sliced for examination of injection sites. Mice with missed injections were either assigned to a separate group or excluded in final data analyses.

### Physiological measurements

In the *Ntrk2* gene deletion experiments, body weight was measured weekly. Body length (naso-anal) was measured from anesthetized mice in the end of each experiment. For food intake, mice were individually housed, and daily food intake was measured over 5 d after 3 d of acclimation. Body composition was determined using a Minispec LF-50/mq 7.5 NMR Analyzer (Brucker Optics). VO_2_ and locomotor activity were assessed with a comprehensive lab animal monitoring system (CLAMS, Columbus Instruments). Ambulatory activity was measured as light beam breaks in the XY horizontal plane.

### Chemogenetic manipulation of food intake

Mice injected with AAV2-hSyn-DIO-mCherry, AAV2-hSyn-DIO-hM3D(Gq)-mCherry or AAV2-hSyn-DIO-hM4D(Gi)-mCherry were singly housed for at least 2 weeks with daily handling prior to experiment. For food intake measurement during light cycle, mice were transferred to a new cage without food at 7:30 am and injected intraperitoneally with vehicle or CNO (Sigma-Aldrich #C08325) at 8:30 am. Pre-weighed food was then added to each cage at 9:00 am and food intake was measured hourly for 3 h. For food intake measurement during dark cycle, mice were transferred to a new cage without food at 3 pm and injected with vehicle or CNO at 5:30 pm. Pre-weighed food was then added to each cage at 6:00 pm and food intake was measured hourly from 7:00 pm to 9:00 pm. For food intake measurement after overnight fasting, mice were transferred to a new cage with alpha-dri bedding without food at 6:00 pm for overnight fasting. On the following day, mice were injected with vehicle or CNO at 9:30 am and pre-weighed food was added to each cage at 10:00 am and food intake was measured hourly for 3 h. The CNO dosage was 1 mg/kg body weight for mice injected with AAV2-hSyn-DIO-hM3D(Gq)-mCherry and 3 mg/kg body weight for mice injected with AAV2-hSyn-DIO-hM4D(Gi)-mCherry.

### In situ hybridization

Freshly dissected mouse brains were quickly frozen in a 2-methylbutane (Fisher Scientific #O3551-4) dry ice bath. Subsequently, 20-μm thick cryostat sections were collected on Superfrost Plus slides (Fisherbrand #12-550-15) and used for in situ hybridization. Radioactive in situ hybridization of brain sections was performed using ^35^S-labeled riboprobes as previously described^11^. To generate antisense riboprobes, the mouse cDNA sequence for *Ntrk2* (GenBank accession number X17647, nucleotides 188-722) was amplified by PCR and cloned into pBluscript II KS (-) plasmid (Stratagene). Antisense probes were synthesized with T3 RNA polymerase (Promega #2083) and labeled with alpha-^35^S-CTP (PerkinElmer #NEG064H250UC) and alpha-^35^S-UTP (PerkinElmer #NEG039H250UC). Brain sections were postfixed in 4% paraformaldehyde for 1 h and acetylated for 10 min with 0.1M triethanolamine hydrochloride/0.25% acetic anhydride (pH 8.0). After dehydration, hybridization was performed with ^35^S labeled probes at 55°C for overnight. After hybridization, sections were treated with RNase A (200 μg/ml, Fisher Scientific # BP2539100) to remove unannealed probes. After dehydration, sections were exposed to Kodak BioMax MR films (Carestream Health) for 3 d. Developed film was scanned at 1,200 dpi for image processing.

### Statistical analysis

All data are expressed as mean ± SEM. Data were tested using Student’s *t* test or two-way ANOVA with post hoc Bonferroni’s or Tukey’s multiple comparisons for statistical significance.

