## Supplementary Information for "TrkB-expressing paraventricular hypothalamic neurons suppress appetite through multiple neurocircuits"

**Supplementary Figures**

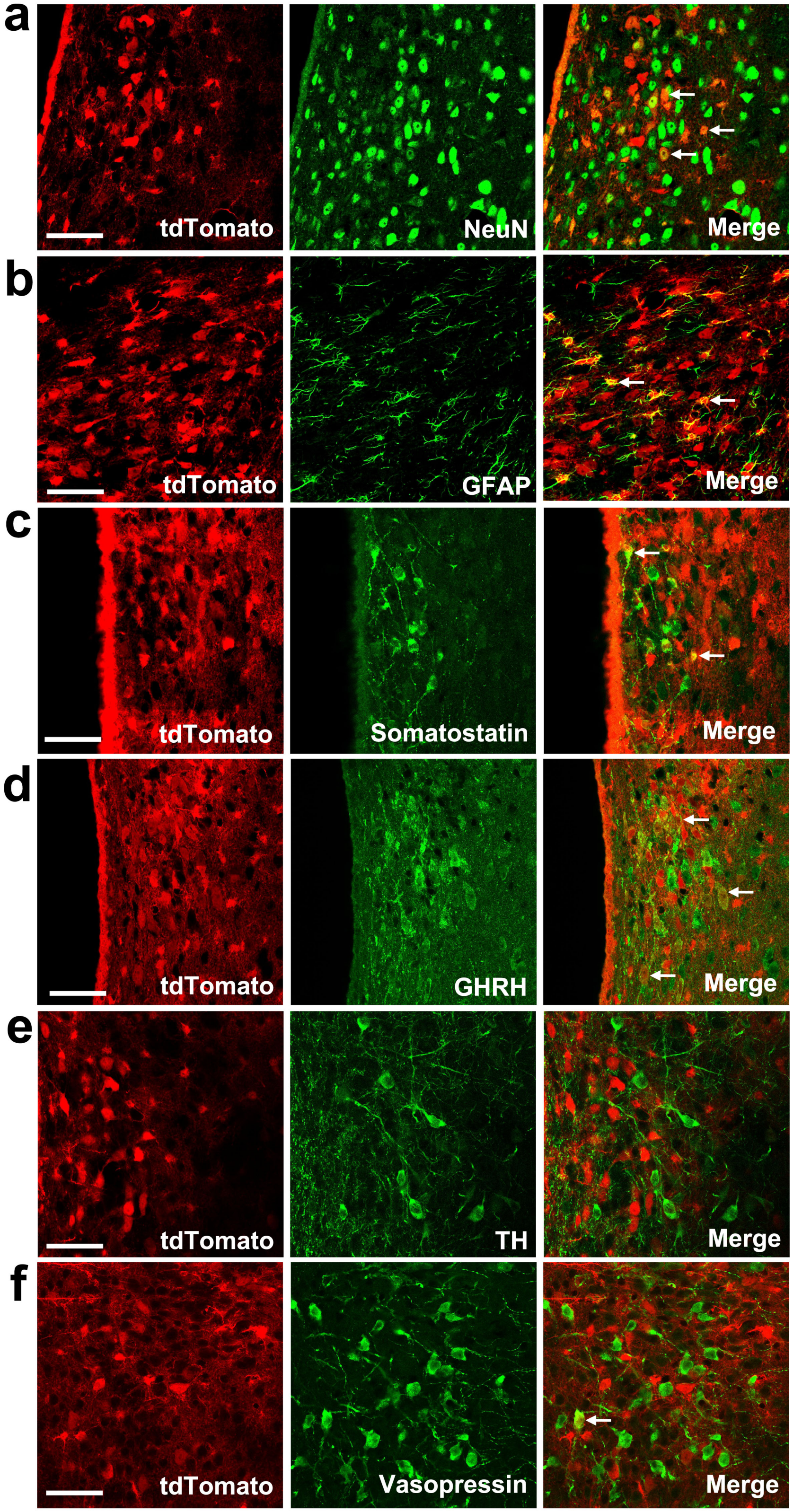

**Supplementary Figure 1. Co-expression of TrkB with cellular markers in the PVH.** TrkB-expressing cells are marked by tdTomato in tamoxifen-treated *Ntrk2*$^{CreER/+}$*;Rosa26*$^{Ai9/+}$ mice. (**a & b**) The majority of TrkB-expressing cells in the PVH are neurons marked by NeuN, while some of TrkB-expressing cells are GFAP-expressing astrocytes. (**c & d**) Some PVH$^{TrkB}$ neurons express either somatostatin or GHRH. (**e & f**) Very few PVH$^{TrkB}$ neurons express tyrosine hydroxylase (TH) or vasopressin. Arrows denote representative neurons expressing TrkB and another cellular marker. Scale bars are 50 μm long.

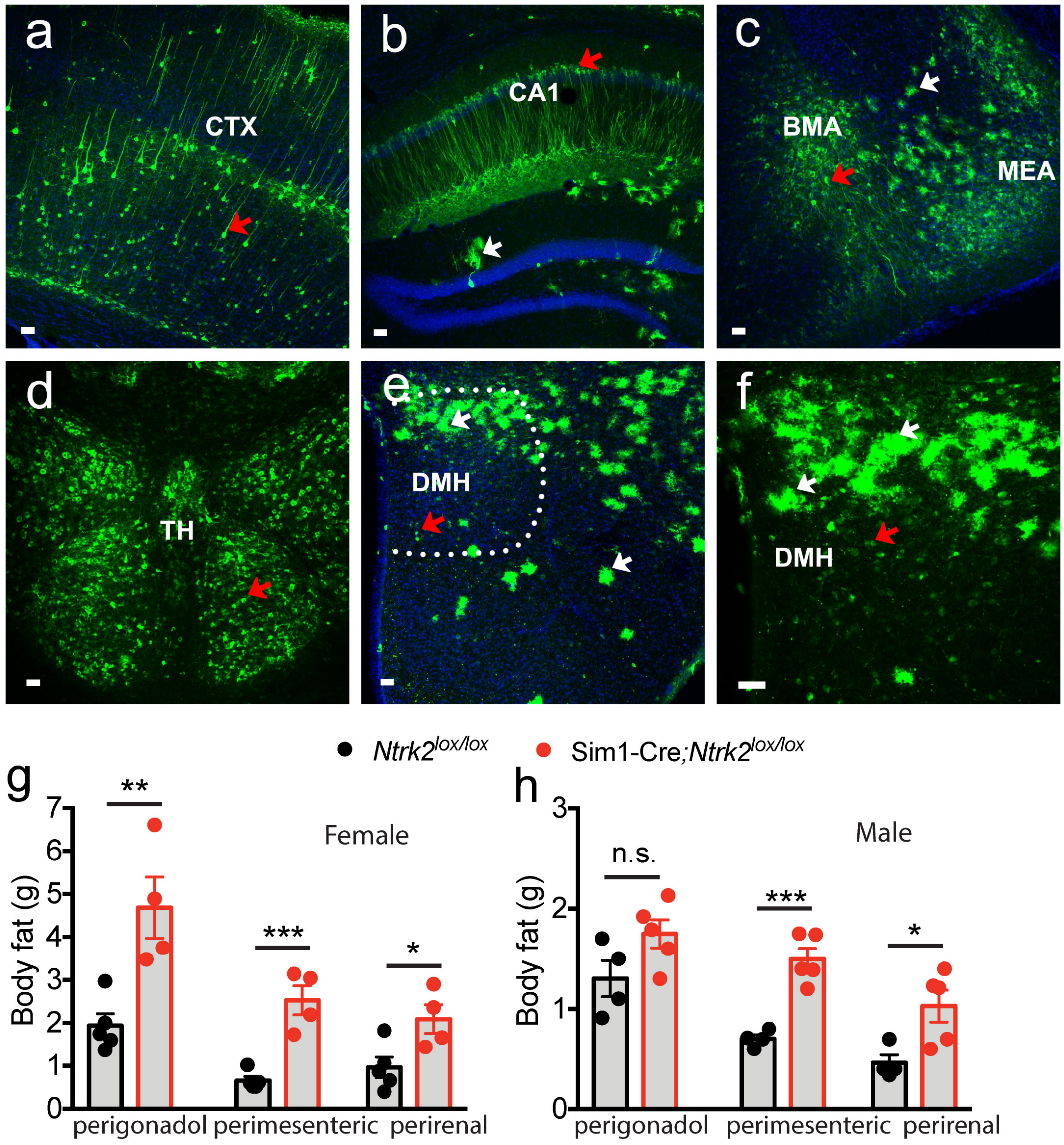


**Supplementary Figure 2. Deletion of the *Ntrk2* gene using the Sim1-Cre transgene.** (**a-f**) β-galactosidase immunohistochemistry of Sim1-Cre;*Ntrk2*$^{fBZ/+}$ brain sections, showing abolishment of the *Ntrk2* expression in some cells in the cerebral cortex (a), hippocampal CA1 region (b), amygdala (c), thalamus (d), and DMH (e & f). Red and white arrows denote β-galactosidase-expressing neurons and astrocytes, respectively. CTX, cerebral cortex; BMA, basomedial amygdala; MEA, medial amygdala; TH, thalamus; DMH, dorsomedial hypothalamus. Scale bars are 50 μm long. (**g & h**) Weight of perigonadal, perimesenteric and perirenal adipose tissues in 16-wk-old female and male *Ntrk2*$^{lox/lox}$ (control) and Sim1-Cre;*Ntrk2*$^{lox/lox}$ (mutant) mice. n = 5 controls and 4 mutants for each sex. Student's *t* test; *$P < 0.05$, **$P < 0.01$, and ***$P < 0.001$.

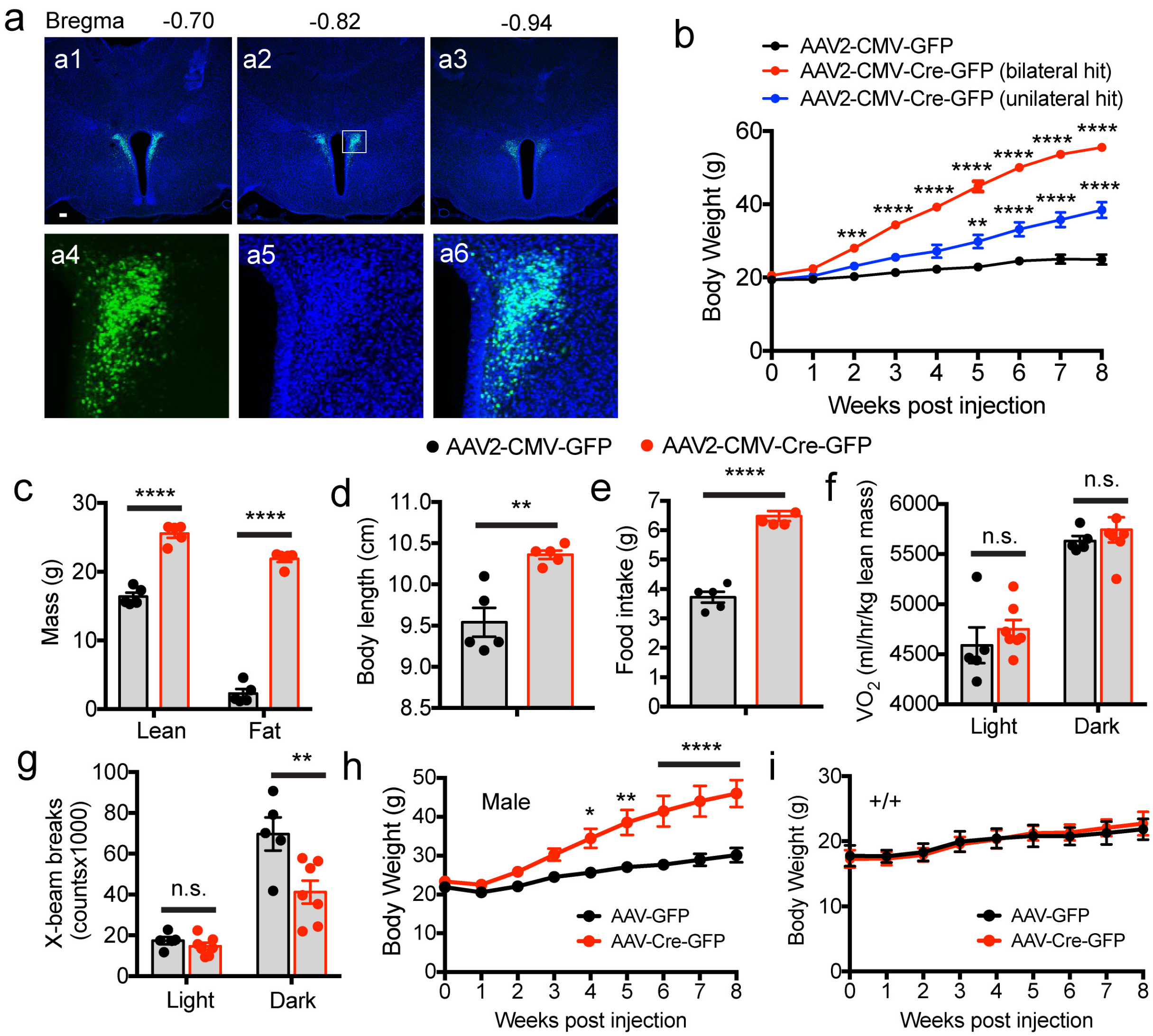


**Supplementary Figure 3. Deletion of the *Ntrk2* gene in the PVH of adult mice leads to hyperphagic obesity.** (**a**) Confocal images showing bilateral injection of AAV2-CMV-Cre-GFP into the PVH of female *Ntrk2*$^{lox/lox}$ mice (a1-a3) and intactness of the PVH after 9 weeks of AAV transduction (a4 – a6). Scale bar is 50 μm long. (**b**) Body weight of female *Ntrk2*$^{lox/lox}$ mice injected with either AAV2-CMV-GFP or AAV2-CMV-Cre-GFP (100 nl) into the PVH bilaterally. Mice injected with AAV2-CMV-Cre-GFP are divided into two groups on the basis of either bilateral or unilateral hit. n = 5 mice for each group. Two-way ANOVA with post hoc Bonferroni multiple comparisons; $F_{(2, 108)} = 488.0$, $P < 0.0001$ for viral injection; **$P < 0.01$, ***$P < 0.001$ and ****$P < 0.0001$ when compared with the AAV2-CMV-GFP group. (**c**) Body composition of female *Ntrk2*$^{lox/lox}$ mice 8 weeks after AAV injection. Student's *t* test; ****$P < 0.0001$. (**d**) Body length of female *Ntrk2*$^{lox/lox}$ mice 8 weeks after AAV injection. Student's *t* test; **$P < 0.01$. (**e**) Daily food intake of female *Ntrk2*$^{lox/lox}$ mice during post-injection week 4. Student's *t* test; ****$P < 0.0001$. (**f**) $O_2$ consumption of female *Ntrk2*$^{lox/lox}$ mice during post-injection week 4. Two-way ANOVA with post hoc Bonferroni multiple comparisons; n.s., not significant. (**g**) Locomotor activity of female *Ntrk2*$^{lox/lox}$ mice during post-injection week 4. Two-way ANOVA with post hoc Bonferroni multiple comparisons; n.s. = not significant and **$P < 0.01$. (**h**) Body weight of male *Ntrk2*$^{lox/lox}$ mice injected with either AAV2-CMV-GFP or AAV2-CMV-Cre-GFP into the PVH bilaterally. n = 6 mice for AAV2-CMV-GFP and 7 mice for AAV2-CMV-Cre-GFP. Two-way ANOVA with post hoc Bonferroni multiple comparisons; $F_{(1, 90)} = 80.53$, $P < 0.0001$ for viral injection; *$P < 0.05$, **$P < 0.01$, and ***$P < 0.0001$. (**i**) Body weight of female WT mice injected with either AAV2-CMV-GFP or AAV2-CMV-Cre-GFP into the PVH bilaterally. n = 3 mice for AAV2-CMV-GFP and 7 mice for AAV2-CMV-Cre-GFP. Two-way ANOVA with post hoc Bonferroni multiple comparisons; $F_{(1, 64)} = 0.01326$, $P = 0.9112$ for viral injection. Error bars indicate SEM.

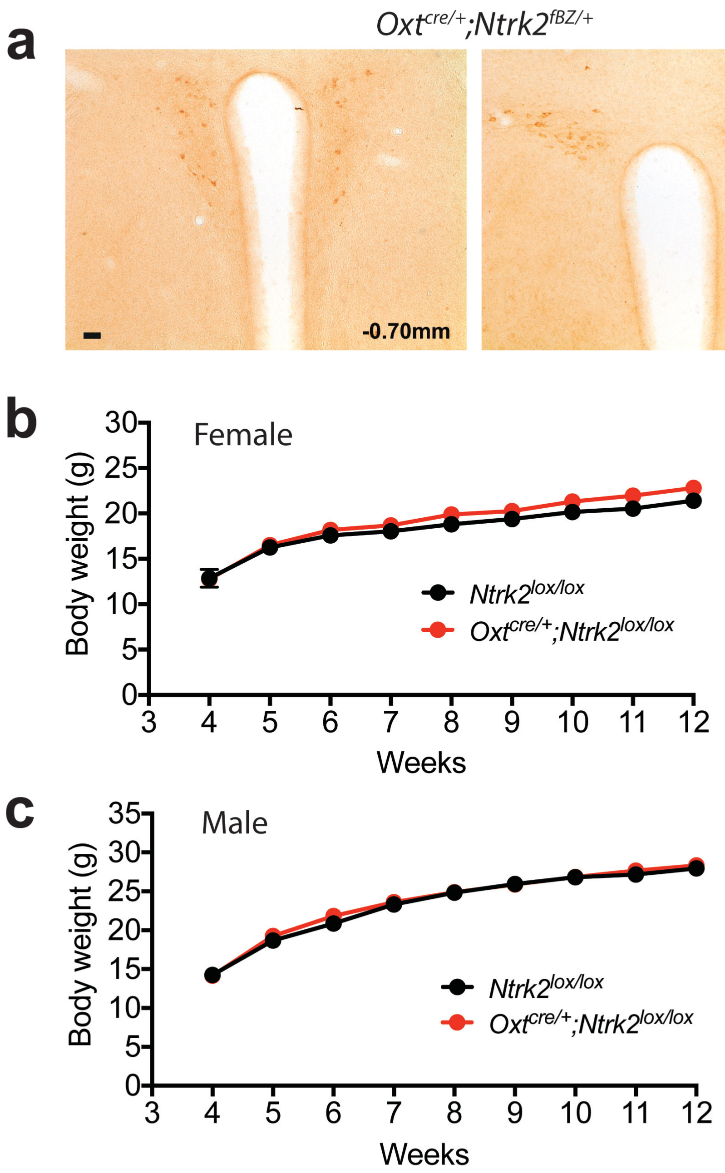


**Supplementary Figure 4. Deletion of the *Ntrk2* gene in oxytocin-expressing cells does not alter body weight.** (**a**) β-galactosidase immunohistochemistry on $Oxt^{Cre/+}$;$Ntrk2^{fBZ/+}$ brain sections. Positive cells indicate abolishment of *Ntrk2* expression. The approximate locations of the two brain sections relative to the bregma are indicated. Scale bar is 50 μm long. (**b**) Body weight of female $Ntrk2^{lox/lox}$ and $Oxt^{Cre/+}$;$Ntrk2^{lox/lox}$ mice. n = 6-8 mice per genotype. Two-way ANOVA with post hoc Bonferroni multiple comparisons; $F_{(1, 96)} = 1.482$, $P = 0.2468$ for genotype. (**c**) Body weight of male $Ntrk2^{lox/lox}$ and $Oxt^{Cre/+}$;$Ntrk2^{lox/lox}$ mice. n = 8-9 mice per genotype. Two-way ANOVA with post hoc Bonferroni multiple comparisons; $F_{(1, 120)} = 3.256$, $P = 0.6477$ for genotype.

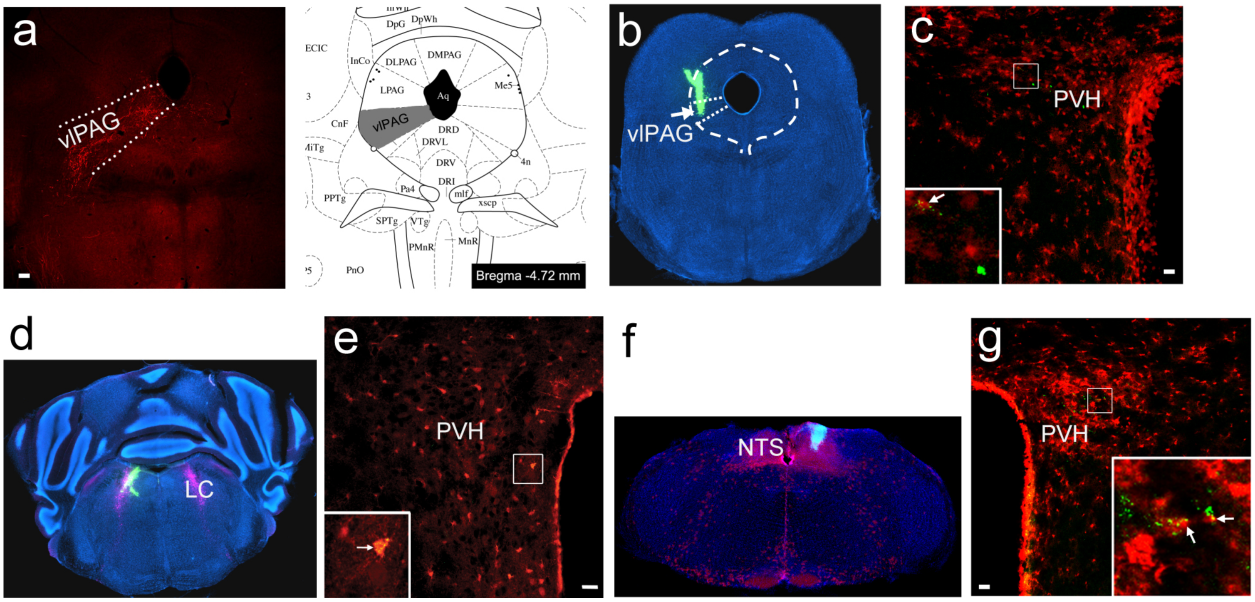


**Supplementary Figure 5. Projections of PVH$^{TrkB}$ neurons.** (**a**) AAV2-CAG-FLEX-tdTomato was injected into the PVH of *Ntrk2$^{CreER/+}$* mice. After the mice were treated with tamoxifen, tdTomato-labeled axonal terminals were detected in the ventrolateral periaqueductal gray (vlPAG). (**b & c**) Green retrobeads (GRBs) were unilaterally injected into the vlPAG of tamoxifen-treated *Ntrk2$^{CreER/+}$;Rosa26$^{Ai9/+}$* mice (b), and GRBs were detected in some tdTomato-labeled PVH$^{TrkB}$ neurons (c). (**d & e**) GRBs were unilaterally injected into the locus coeruleus (LC) of tamoxifen-treated *Ntrk2$^{CreER/+}$;Rosa26$^{Ai9/+}$* mice (d), and GRBs were detected in some PVH$^{TrkB}$ neurons (e). The LC is marked by tyrosine hydroxylase immunoreactivity (pink). (**f & g**) GRBs were unilaterally injected into the nucleus tractus solitaris (NTS) of tamoxifen-treated *Ntrk2$^{CreER/+}$;Rosa26$^{Ai9/+}$* mice (f), and GRBs were detected in some PVH$^{TrkB}$ neurons (g). The scale bar in (a) represents 100 μm and in other panels 50 μm.

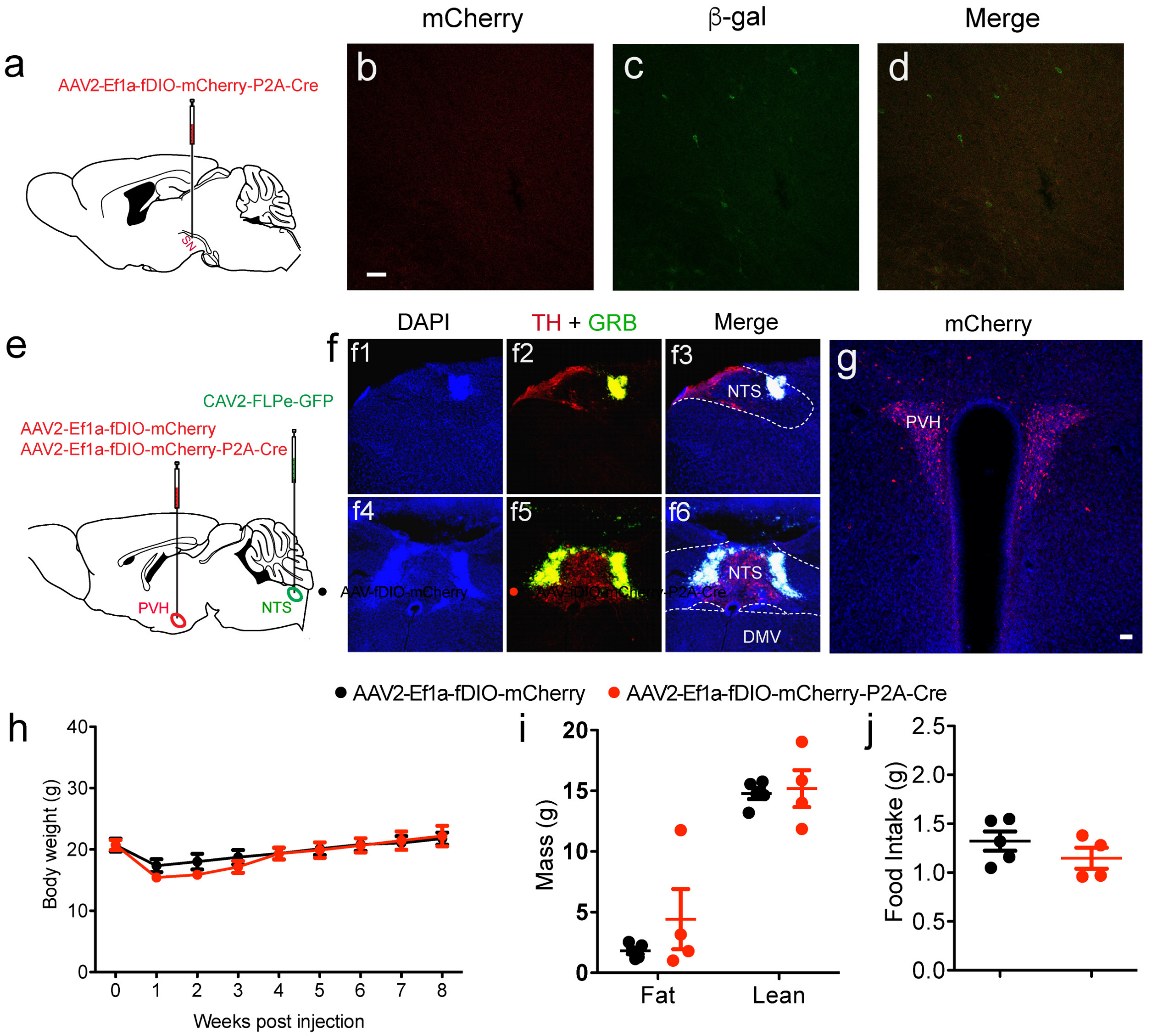


**Figure 6**. Deletion of the *Ntrk2* gene in PVH neurons projecting to the NTS. (**a-d**) Injection of AAV2-Ef1a-fDIO-mCherry-P2A-Cre (200 nl) alone into the substantia nigra did not lead to expression of mCherry and β-galactosidase (β-gal). (**e**) Deletion of the *Ntrk2* gene in PVH neurons projecting to the NTS. A mixture of CAV2-FLPe + GRB (4:1 ratio) and either AAV2-Ef1a-fDIO-mCherry (control) or AAV2-Ef1a-fDIO-mCherry-P2A-Cre (150 nl per site) were bilaterally injected into the NTS and PVH of $Ntrk2^{lox/lox}$ mice, respectively. Co-injected GRBs were used to mark CAV2 injection sites. The CAV2-FLPe + GRB mixture was injected into two sites for each NTS, a rostral site and a caudal site in the central NTS (100 nl per site). (**f**) CAV2 injection sites in the rostral part of central NTS (f1-3) and the caudal part of central NTS (f4-6). Tyrosine hydroxylase (TH) immunoreactivity (red) outlines the NTS, while GRBs (green) indicate CAV2 injection sites. (**g**) Neurons that were transduced by both CAV2-FLPe and AAV2-Ef1a-fDIO-mCherry-P2A-Cre expressed mCherry in the PVH. The image is a projection of several z optical sections. (**h-j**) Body weight (h), fat mass and lean mass (i), and 4-hour food intake (j) of female $Ntrk2^{lox/lox}$ mice injected with CAV2-FLPe into the NTS and either AAV2-Ef1a-fDIO-mCherry or AAV2-Ef1a-fDIO-mCherry-P2A-Cre into the PVH. Food intake was measured in the first 4 hours of the dark cycle. Body weight data were analyzed using two-way ANOVA with Bonferroni *post hoc* tests; $F_{(1, 96)} = 0.1720$,

*P* = 0.6856 for virus (n=6 mice for control and 8 mice for Cre). Student's *t* test reveals no significant difference in body composition and food intake between groups (n=5 mice for control and 4 mice for Cre). Scale bars represent 50 μm.

### Supplementary Table 1. Infection scores for AAV-Cre-GFP injected female *Ntrk2*$^{lox/lox}$ mice

| | | | Female AAV-Cre Bilateral PVH Scores & Non-PVH Total Scores | | | | | | | | | | |
|---|---|---|---|---|---|---|---|---|---|---|---|---|---|
| | | | Missed | | | | | Hit | | | | | |
| PVH | Anterior | **dorsal:** | + | - | + | - | + | - | + | - | ++ | ++++ | ++++ |
| | | **ventral:** | + | - | - | - | ++ | - | - | - | - | +++ | +++ |
| | Central | **dorsomedial:** | - | - | - | + | - | + | - | ++++ | ++++ | ++++ | +++++ |
| | | **dorsolateral:** | - | - | + | + | - | + | +++ | ++++ | +++++ | ++++ | +++++ |
| | | **ventral:** | - | - | - | - | + | ++ | + | +++ | ++ | +++ | ++++ |
| | Posterior | **medial:** | - | + | - | + | - | +++ | ++ | ++++ | ++ | +++ | +++ |
| | | **lateral:** | - | + | - | - | - | + | + | + | + | + | + |
| | | **Overall Bilateral PVH Score:** | **2** | **2** | **2** | **3** | **4** | **8** | **8** | **16** | **16** | **22** | **25** |

| | | | | | | | | | | | | | |
|---|---|---|---|---|---|---|---|---|---|---|---|---|---|
| Non-PVH Hypothalamus | | Anterior (AH): | - | - | - | - | •• | • | • | • | - | - | - |
| | | Dorsmedial (DMH): | - | • | - | - | - | • | - | • | - | - | - |
| | | Ventromedial (VMH): | - | • | - | - | - | - | • | • | - | - | - |
| | | Lateral (LH): | - | - | - | - | - | - | - | - | - | - | - |
| | | Arcuate (ARC): | - | - | - | - | - | - | - | - | - | - | - |
| | | Posterior (PH): | - | •• | - | •• | - | - | - | - | - | - | • |
| | | Suprachiasmatic (SCN): | - | - | - | - | - | - | - | - | - | - | - |
| | | Medial Preoptic (MPO): | •• | - | - | - | •• | - | • | - | - | - | - |
| | | Median Preoptic (MnPO): | • | - | - | - | - | - | - | - | - | - | - |
| | | Zona Incerta: | - | • | • | - | • | • | • | • | • | • | • |
| | | Subincertal Nucleus: | - | - | - | • | - | • | - | - | • | - | • |
| | | Bed Nucleus of Stria Terminalis: | • | - | - | - | • | - | - | - | - | - | - |
| THALAMUS | | Mediodorsal (MD): | • | ••• | •• | •••• | - | ••• | •• | •• | •• | •• | - |
| | | Paratenial (PT): | •••• | - | •• | - | •• | •• | - | - | •• | •• | - |
| | | Submedius (Sub): | - | • | - | •• | - | • | - | •• | • | •• | - |
| | | Reuniens (Re): | - | • | - | •• | - | - | • | • | • | • | • |
| | | Anteromedial (AM): | - | - | - | ••• | - | •• | •• | - | - | - | - |
| | | Paraventricular (PVT): | •• | - | - | - | •• | •• | • | - | - | - | - |
| | | Interanteromedial (IAM): | - | - | - | • | - | - | - | - | • | • | •• |
| | | Rhomboid (Rh): | - | - | - | ••• | - | - | - | - | - | - | - |
| | | Central Medial (CM): | - | - | - | •• | - | - | - | - | - | - | - |
| | | Interanterodorsal (IAD): | - | - | - | - | - | - | - | - | - | - | • |
| | | Ventromedial (VM): | - | - | - | • | - | - | - | - | - | - | - |
| | | Habenula: | - | • | - | • | - | - | - | - | - | - | - |
| | | Hippocampus: | - | • | - | • | - | - | - | - | - | - | - |
| | | Cingulate/Motor Cortex: | - | • | - | • | • | - | • | - | - | - | - |
